# AQP1 Differentially Orchestrates Endothelial Cell Senescence

**DOI:** 10.1101/2024.03.13.584782

**Authors:** Khatereh Shabanian, Taraneh Shabanian, Gergely Karsai, Sandra Lettlova, Luca Pontiggia, Frank Ruschitzka, Jürg H. Beer, Seyed Soheil Saeedi Saravi

## Abstract

Accumulation of senescent endothelial cells (ECs) with age is a pivotal driver of cardiovascular diseases in aging. However, little is known about the mechanisms and signaling pathways that regulate EC senescence. In this report, we delineate a previously unrecognized role of aquaporin 1 (AQP1) in orchestrating extracellular hydrogen peroxide (H_2_O_2_)-induced cellular senescence in aortic ECs. Our findings underscore AQP1’s differential impact on senescence hallmarks, including cell-cycle arrest, senescence-associated secretory phenotype (SASP), and DNA damage responses, intricately regulating angiogenesis. In proliferating ECs, AQP1 is crucial for maintaining angiogenic capacity, whereas disruption of AQP1 induces morphological and mitochondrial alterations, culminating in senescence and impaired angiogenesis. Conversely, *Aqp1* knockdown or selective blockade of AQP1 in senescent ECs rescues the excess H_2_O_2_-induced cellular senescence phenotype and metabolic dysfunction, thereby ameliorating intrinsic angiogenic incompetence. Mechanistically, AQP1 facilitates H_2_O_2_ transmembrane transport, exacerbating oxidant-sensitive kinases CaMKII-AMPK. This process suppresses HDAC4 translocation, consequently de-repressing Mef2A-eNOS signaling in proliferating ECs. However, in senescent ECs, AQP1 overexpression is linked to preserved HDAC4-Mef2A complex and downregulation of eNOS signaling. Together, our studies identify AQP1 as a novel epigenetic regulator of HDAC4-Mef2A-dependent EC senescence and angiogenic potential, highlighting its potential as a therapeutic target for antagonizing age-related cardiovascular diseases.

**Highlights:** • AQP1 is upregulated in aortic endothelial cells with aging

• AQP1 differentially orchestrates H_2_O_2_-mediated EC senescence

• AQP1 plays a dual role in regulating angiogenesis in proliferating and senescent ECs

• AQP1 controls EC function by differentially modulating HDAC4-Mef2A pathway

• AQP1 deficiency restores angiogenic capacity in senescent ECs

## Introduction

Aging, a significant risk factor for cardiovascular diseases, is characterized by cellular senescence [1,2]. This phenomenon is a permanent cell cycle arrest linked to accumulated stochastic cellular damage [3,4]. Cellular senescence, particularly evident in vascular endothelial cells (ECs), contributes to cardiovascular pathologies like atherosclerosis and hypertension in aging [5,6]. Understanding the triggers and mechanisms driving EC senescence holds promise for developing novel therapies to clear senescent cells and subsequently ameliorate the age-related pathologies and extend healthspan [7]. Cellular senescence is orchestrated through a variety of networks triggered by extracellular and intracellular stimuli. Elucidating the senescence regulatory mechanism could provide potential intervention targets to attenuate degenerative processes and improve the quality of life for the aging population. Reactive oxygen species (ROS) are the stimuli that induce oxidative stress which is closely associated with cellular senescence [8]. Existing evidence collectively suggests that oxidative stress can induce senescence-like features in ECs by promoting both excessive ROS accumulation and disrupting their redox state [1]. Oxidative stress is primarily involved in the oxidation of all essential macromolecules and metabolic disorders, ultimately leading to impaired energy homeostasis and cell cycle arrest in ECs [].

Hydrogen peroxide (H_2_O_2_), a stable ROS, plays a dual role in ECs, acting as both a signaling molecule involving in oxidative stress and a physiological effector depending upon its concentration [10]. The fluctuating H_2_O_2_ levels in the extracellular and intracellular space are crucial for cellular metabolism and cell cycle [11]. Aging is associated with an increase in H_2_O_2_ levels in both extracellular space and intracellular compartments, with its extracellular production occurring through various enzymatic and non-enzymatic reactions [12,13]. Identifying key regulators responsible for H_2_O_2_ transmembrane transport and its distribution into the intracellular space is paramount for designing targeted senotherapies. While traditionally considered freely diffusible across plasma membranes, recent studies suggest a vital role for aquaporin (AQP) channels, also known as peroxiporins, in facilitating H_2_O_2_ diffusion into the cell cytosol [14,15]. Among several AQP isoforms, AQP1, widely expressed in vascular endothelia, regulates not only water but also H_2_O_2_ transport across the plasma membrane of these cells [16–18]. For the past two decades, investigations have revealed that genetic deficiency of AQP1 in AQP1-null mice and primary cultures of aortic endothelial cells leads to significant impairments in angiogenesis [16]. Despite this, whether the involvement of H_2_O_2_ transport regulation has been taken into account in mediating the role of AQP1 in endothelial cell function has remained largely unexplored. In previous studies, it was elucidated that AQP1-mediated transmembrane transport of extracellularly produced H_2_O_2_ is critical for activating hypertrophic signals within cardiac myocytes, subsequently leading to cardiac hypertrophy. Deletion of *Aqp1* or blockade of AQP1 could present a promising avenue for the treatment of hypertrophic cardiomyopathies [14]. On the other hand, the aging process may confer significant effects on the distribution and functionality of AQP1 across various cell types and tissues. Notably, within cardiac tissue, AQP1 upregulation has been identified as a compensatory mechanism to address water and electrolyte imbalances associated with aging [19,20]. However, our understanding of age-related alterations in AQP1 expression and function within endothelial cells remains limited, let alone its involvement in regulating H_2_O_2_ levels in EC senescence.

In this work, we conducted a comparative analysis of AQP1 expression in aortic ECs derived from young and aged mice, revealing an age-associated upregulation of AQP1 concurrent with endothelial dysfunction. Further study unveiled the pivotal role of AQP1 in modulating EC senescence through the regulation of H_2_O_2_ transmembrane transport. Specifically, we observed that in proliferating ECs (PECs), AQP1 is necessary for HDAC4-mediated activation of Mef2A-eNOS signaling, promoting angiogenesis. Conversely, in senescent ECs (SECs), AQP1 impeded HDAC4 nuclear export, resulting in the downregulation of Mef2A and eNOS phosphorylation. Most importantly, targeted manipulation of AQP1 expression via silencing or pharmacological blockade with Bacopaside II effectively reversed EC senescence and restored angiogenic capacity. Together, these findings not only elucidate a previously unexplored mechanism by which AQP1 modulates EC senescence and angiogenesis but also underscore the senotherapeutic potential of AQP1 deficiency in aging-related vascular decline.

## Results

### Aging Leads to Upregulation of AQP1 in Endothelial Cells in Humans and Mice

To examine how aging contributes to alteration of AQP1 expression in endothelial cells, we isolated endothelial cells from aortas of both young (3 months) and old (24 months) mice and examined AQP1 expression by immunobloting (Fig. 1A). Our data identified markedly increased expression of AQP1 in ECs from old mice compared to young mice (Fig. 1B). We then stained aortas for CD31 (biomarker of endothelial cells; cyan) and AQP1 (green) In order to confirm what seen in Fig. 1B. We observed a remarkable abundance of AQP1 in CD31^+^ aortic ECs from old mice as opposed to young mice (Fig. 1C). These findings were accompanied by a markedly reduced endothelial function in old mice, as reflected by decreased endothelium-dependent relaxation to acetylcholine, compared to what seen in young mice. As shown in Supplemental Fig. 1, there was a significantly reduced maximal relaxation at 10^−5^☐M (*p*☐<☐0.001; Young: 98.43☐±☐6.06; Old: 62.06☐±☐8.74), and rightward shift of dose-response curve to acetylcholine (*p*☐<☐0.05; EC_50_, Young: 32.01☐±☐6.23 nM; Old: 51.69☐±☐11.89 nM) was seen in isolated aortas from young *versus* old mice. Additionally, we tested the AQP1 expression in proliferating human aortic endothelial cells (PEC; passage 4-5) and replicative senescent HAECs (SEC; passage 15-17). Of note, senescence phenotype (such as enlarged, flattened, and multinucleated appearance of cells) was verified in SECs (Supplemental Fig. 2). As seen in Fig. 1D,E, cellular senescence, as a hallmark of aging, is associated with significant AQP1 upregulation in SECs as opposed to PECs.

**Fig. 1.**
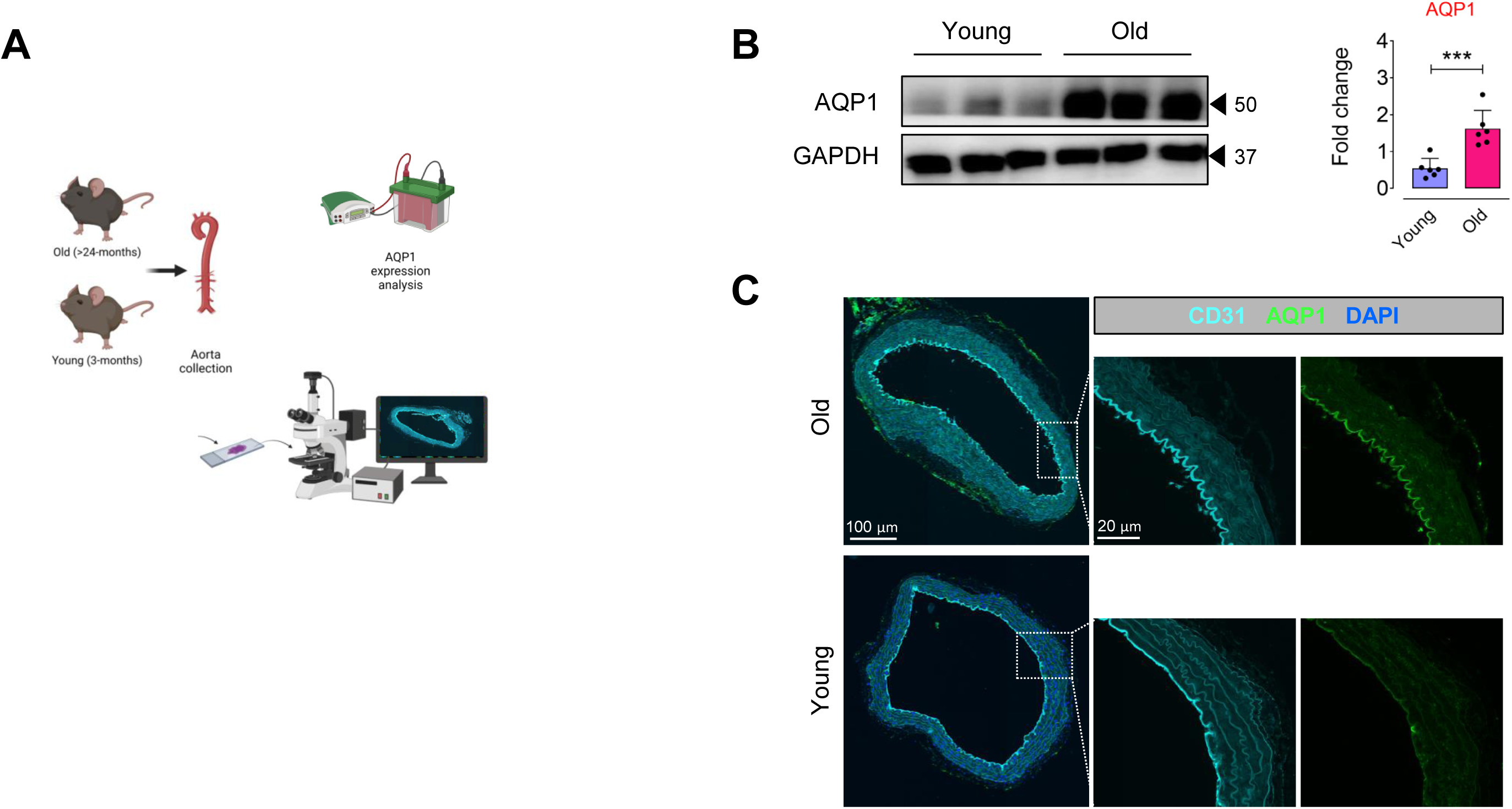
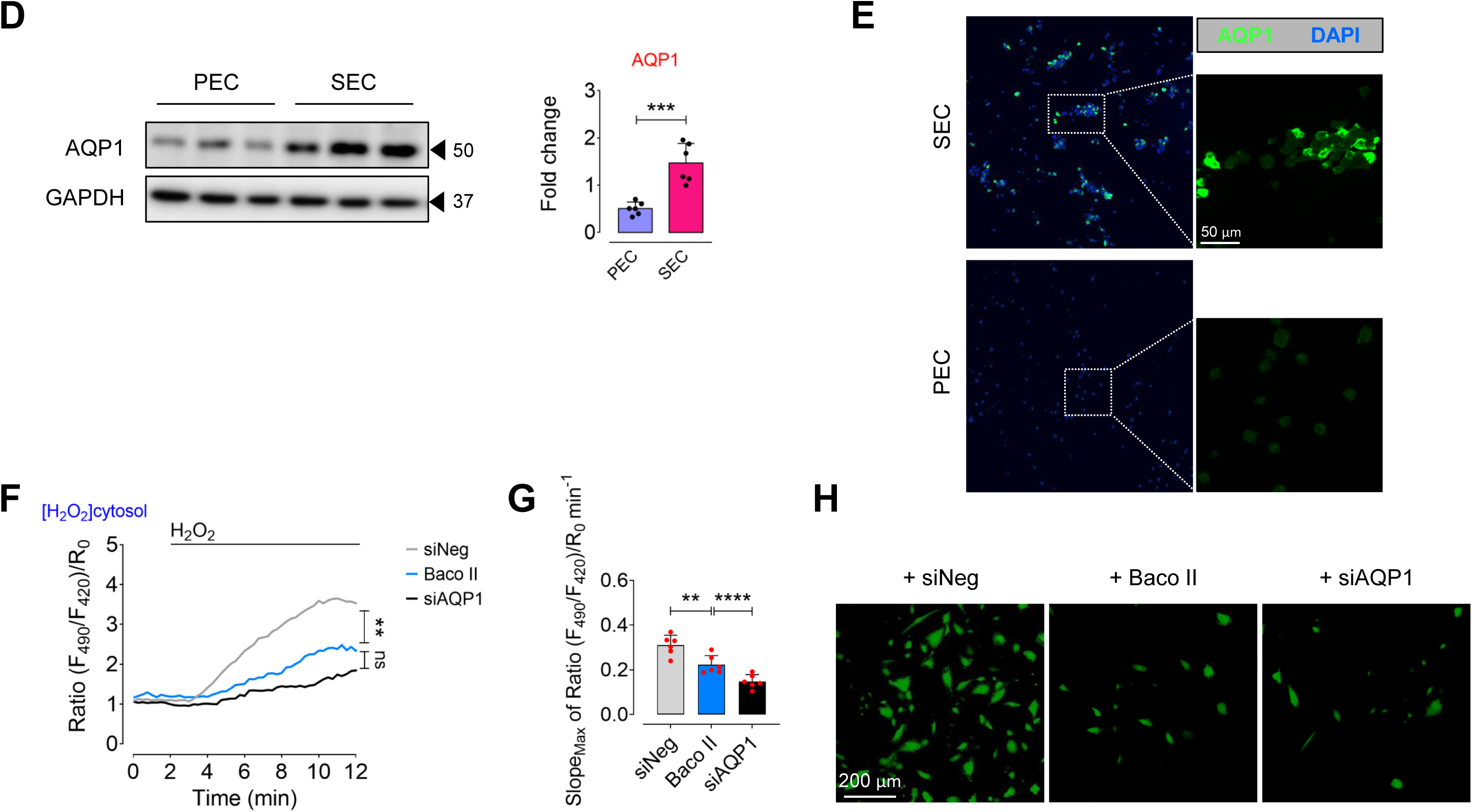
Aging is associated with AQP1 upregulation in aortic endothelial cells. **A**, Schematic diagram of the experimental setting: Aortic ECs were collected from >24 (old) and 3 (young)-month-old C57BL/6J male mice for AQP1 immunoblotting and immunostaining. **B,** Immunoblot analysis demonstrates a significant age-related increase in the expression of AQP1 in aortic ECs of mice (n=6). **C,** AQP1 immunostaining demonstrates higher expression of AQP1 in the ascending aortic endothelial cells from old mice compared to those from young mice (n=5-6). **D,E,** Immunoblots and immunostaining represent a marked higher expression of AQP1 in senescent ECs (SEC; at p15-17) compared to proliferating ECs (PEC; at p4-5) (n=6). **F,** PECs were transduced with adenovirus 5 (AV5)-HyPer7.2 targeted to the cell cytosol for ratiometric fluorescence imaging to detect cytosolic H_2_O_2_ levels in the presence exogeneous H_2_O_2_ (50 μM). The curves demonstrate significantly lower H_2_O_2_ responses in the cytosol of HyPer7.2-transduced PECs following treatment with either Baco II or siAQP1 as compared to that in siNeg-transfected PECs. **G,** Bar chart shows maximal HyPer7.2-ratio changes over time (slope) in the cytosol of PECs exposed to siNeg, Baco II, or siAQP1 (n=6). **H,** CellRox green staining reveals a markedly lower generation of intracellular ROS, including H_2_O_2_, (as green fluorescence) in siNeg-transfected PECs as opposed to that in Baco II-or siAQP1-exposed PECs. Scale bars, 20 and 100 μm. Error bars represent SD (**B,D,G**). Continuous data are presented as mean ± SD. Statistical analysis was performed with a two-tailed unpaired Student’s *t*-test (**B,D**) and one-way ANOVA followed by Tukey’s post *hoc* test (**F,G**). (\*\**P*<0.01, \*\*\**P*<0.001, \*\*\*\**P*<0.0001, ns, not significant). Source data are provided as a Source Data file.

AQP1 has been recognized to transport H_2_O_2_ in specific cell types [13], which may promote premature cellular senescence [21,22]. To prove the role of AQP1 in regulation of H_2_O_2_-mediated EC senescence, we transduced PECs (control) with ultrasensitive H_2_O_2_ biosensor, AV5-HyPer7.2-NES, and detected H_2_O_2_ levels in the cell cytosol which was markedly reduced following siAQP1 transfection or treatment with a clinically approved AQP1 inhibitor, Bacopaside II, in the presence of exogenous H_2_O_2_ (50 μM; Fig. 1F). Moreover, Fig. 1G illustrates that maximal HyPer7.2 ratio changes over time (slope) significantly reduced in siAQP1-(2.09-fold) or Baco II (1.39-fold)-exposed PECs as opposed to that in siNeg-PECs. Consistently, a significantly higher CellRox green fluorescence (higher number of CellRox^+^ cells), which represents an increased cytosolic ROS, was seen in siNeg-transfected PECs (control) *versus* siAQP1-transfected or Baco II-treated PECs in the presence of exogenous H_2_O_2_ (Fig. 1H). These data verify that both siAQP1 and Baco II could strongly disrupt AQP1 function and decrease the extent and velocity of H_2_O_2_ permeability in the endothelial cells.

The positive association between endothelial AQP1 expression with age in mice alongside concurrent endothelial dysfunction, underscores the potential key role of AQP1 in induction of EC senescence and vascular decline in aging.

### AQP1 Differentially Regulates Endothelial Cell Senescence

According to our findings revealing age-related increase in AQP1 in aortic ECs, AQP1 likely play a pivotal role in senescence in these cells. Therefore, we knocked down *Aqp1* gene (with siRNA silencing) or pharmacologically inhibited AQP1 (with Bacopaside II) in the presence of exogenous H_2_O_2_ to test hallmarks of cellular senescence (Fig. 2A). Surprisingly, genetic silencing or pharmacological inhibition of AQP1 exhibited differential effects on cellular senescence in ECs, validating differential role of AQP1 in promoting EC senescence. Accordingly, both siAQP1-transfected and Baco II-treated PECs exhibited multiple premature senescence-like features including increased numbers of SA-β-gal^+^ cells and γ-H2A.X immunofluorescence (Fig. 2B) comparable to what seen in PECs incubated with H_2_O_2_ alone. We next sought to determine whether AQP1 disruption contributes to cell-cycle arrest in PECs. As a result, a concomitant increase in expression of cyclin-dependent kinase (CDK inhibitors) *p16^INK4a^*, *p19^INK4d^*, and *p21^WAF1/Cip1^*at mRNA level was seen in PECs exposed to either siAQP1 or Baco II (Fig. 2C). In addition, using a qPCR-based assay, we detected the expression of various SASP components, suggesting a positive correlation between AQP1 disruption and upregulation of SASP genes, including *Il1*α, *Il1*β, and *Il6* (Fig. 2D). Our VCAM1 immunostaining also demonstrates an increase in VCAM1 expression in PECs subjected to either siAQP1 or Baco II (Fig. 2E). These were consistent with our single-cell Raman spectroscopy findings that reveal a remarkably different intracellular biochemical content in siNeg-PECs as opposed to those in AQP1-deficient PECs (Supplemental Fig. 3A,B). Peaks and shifts in the spectra indicate differences in molecular composition, potentially reflecting variances in protein and lipid profiles, namely inflammatory cytokines, between the cell populations [23,24]. The distinct Raman spectra seen in these two populations of PECs at specific wavelengths represent disparate potential secretome that was visualized by Raman Linear Discriminant Analysis (LDA). The score plots show that data point clouds corresponding to siNeg-and siAQP1-PEC populations were extremely different (Supplemental Fig. 3C).

**Fig. 2.**
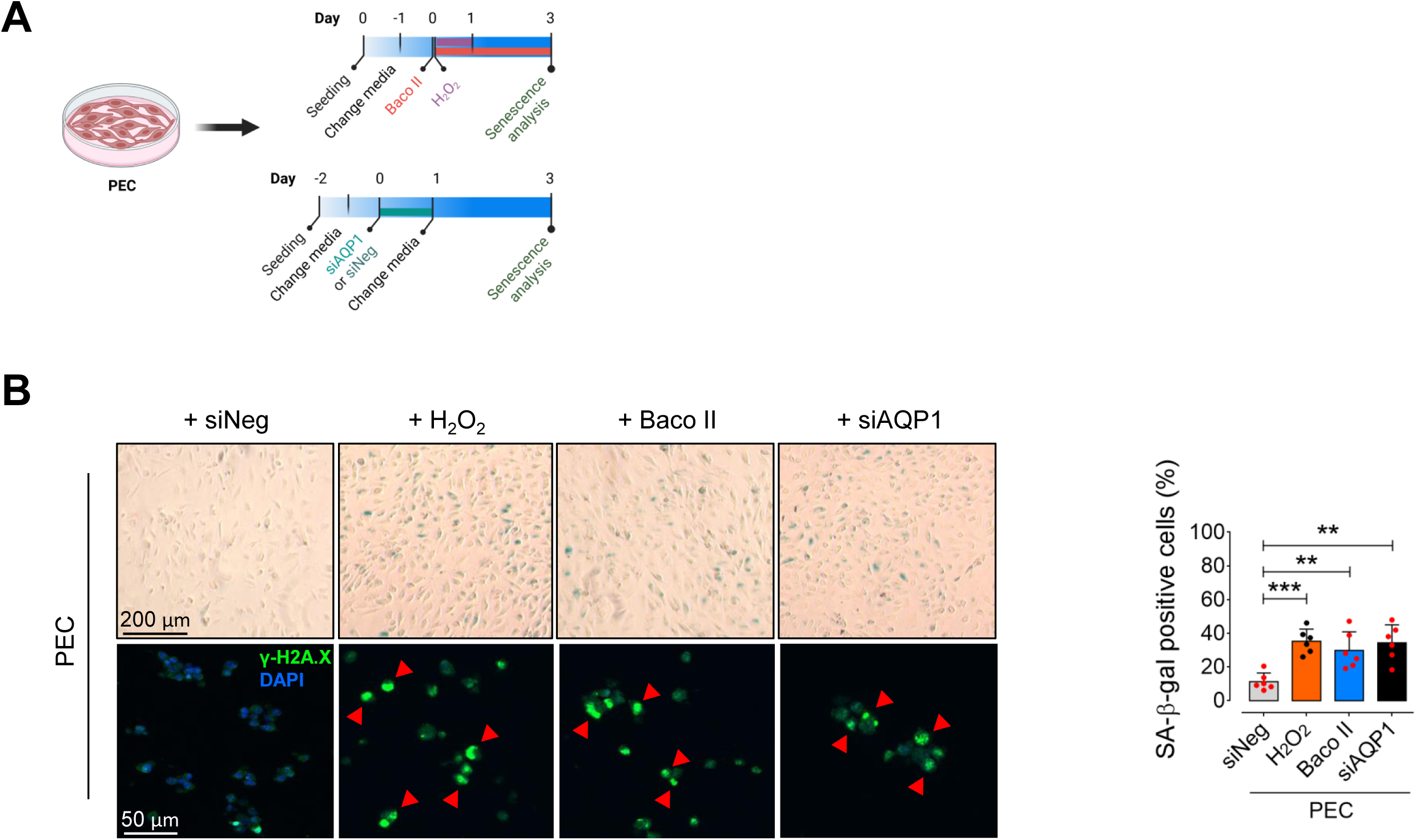
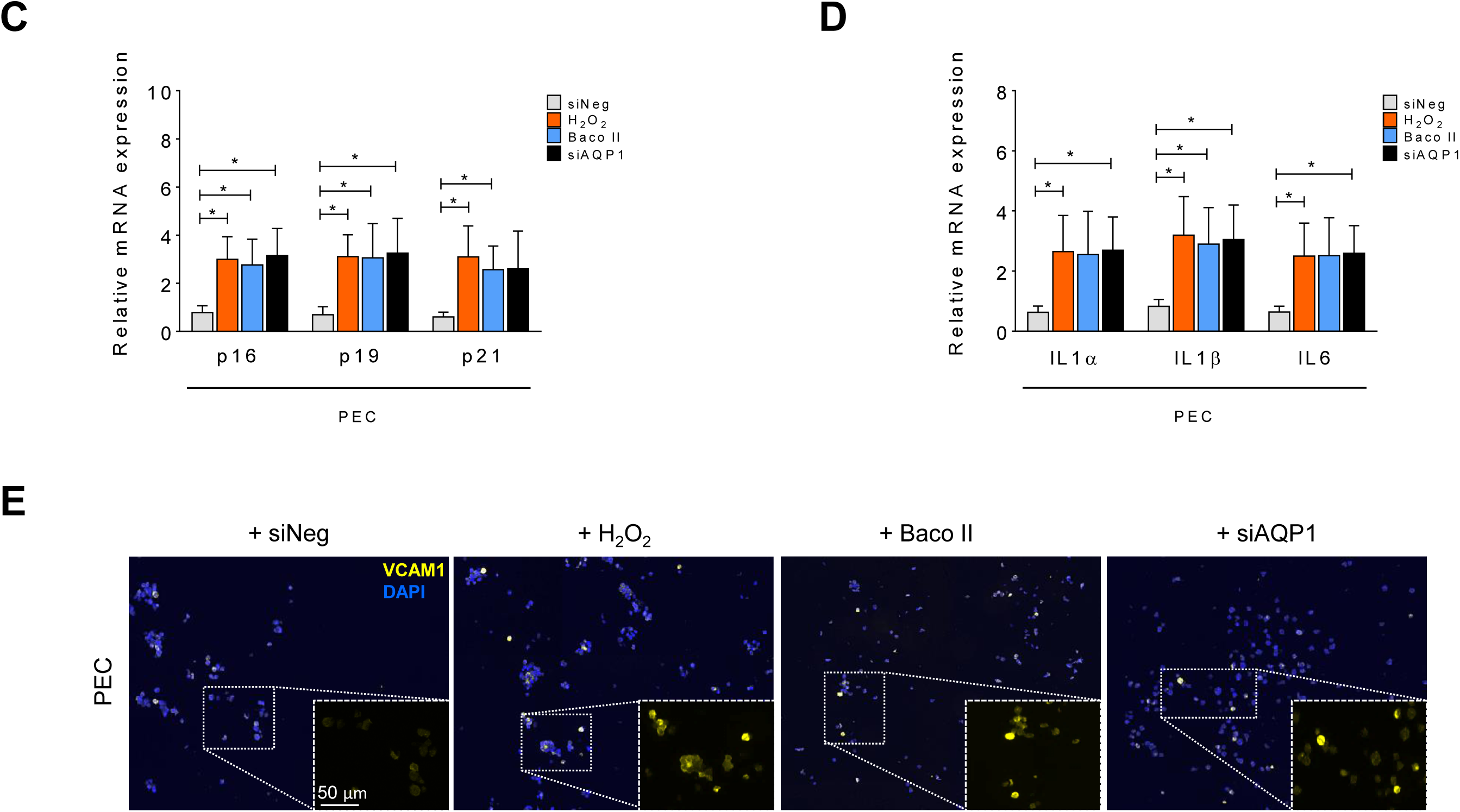
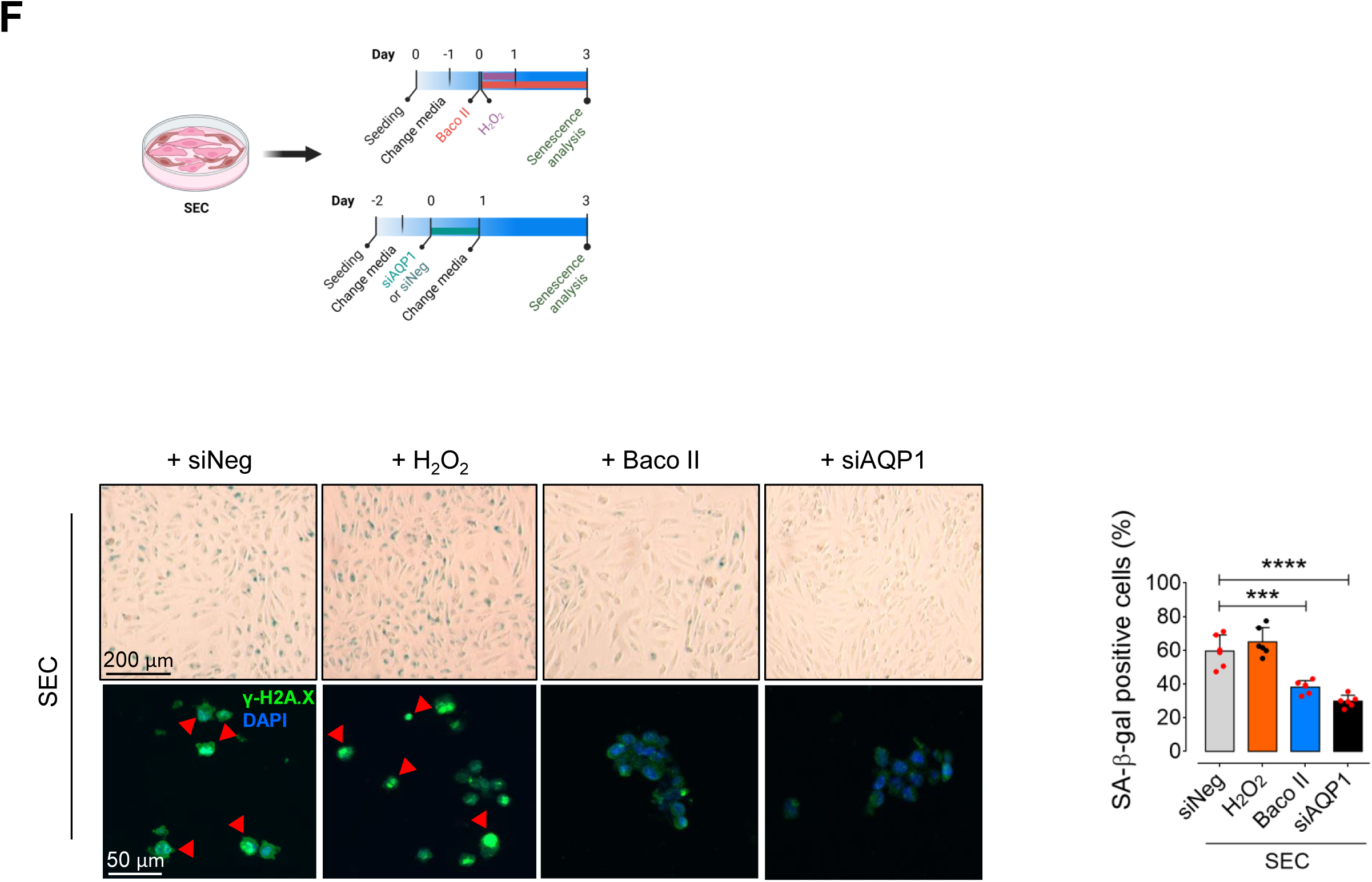
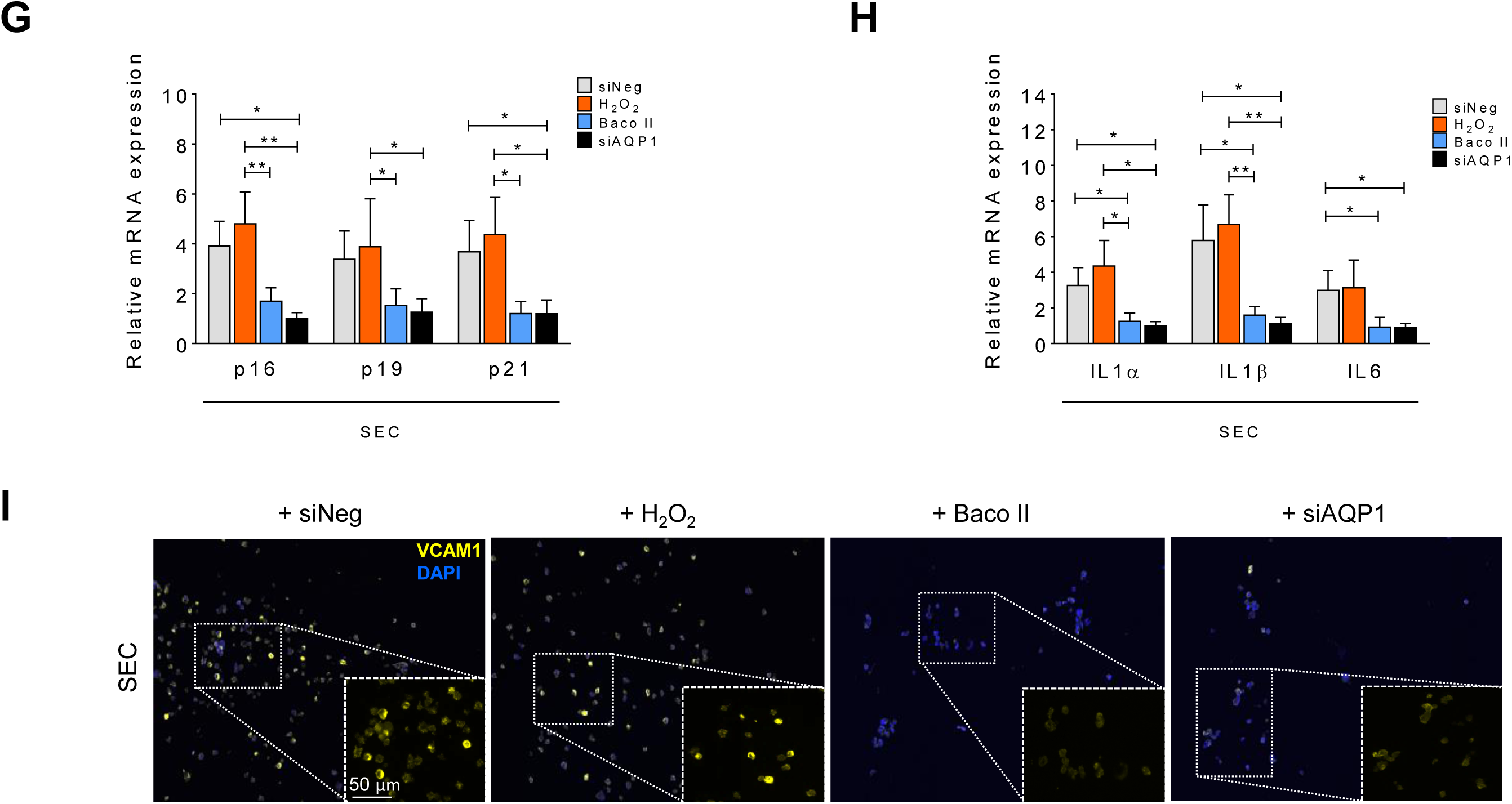
AQP1 differentially regulates endothelial cell senescence. **A**, Schematic diagram of the experimental setting: PECs were treated with exogenous H_2_O_2_ (25 μM) for 24 h or Baco II (10 μM) for 72 h. Another subset of PECs was transfected with siNeg or siAQP1. At day 3, all cells were then subjected to senescence hallmarks profiling. **B,** Left, Upper panel, Representative images show significant increase in the numbers of SA-β-gal^+^ cells in the Baco II-and siAQP1-exposed groups, at the magnitude seen in H_2_O_2_–treated PECs, compared to siNeg-transfected PECs. Right, quantitative plots are shown for SA-β-gal^+^ cells (%) (n=6). Left, Lower panel, γ-H2A.X immunostaining represent DDR in Baco II-and siAQP1-exposed PECs compared to siNeg-transfected PECs (n=6). **C,** qPCR demonstrates that Baco II or siAQP1 treatment resulted in increased expression of CDK inhibitors p16^INK4a^, p19^INK4d^, and p21^WAF1/Cip1^ in PECs. **D,** qPCR shows that SASP components genes IL1α, IL-1β, and IL-6 are significantly upregulated in Baco II-and siAQP1-exposed PECs (n=6). **E,** VCAM1 immunostaining shows its marked overexpression in Baco II-and siAQP1-exposed PECs compared to siNeg-transfected PECs (n=6). **F,** Schematic diagram of the experimental setting: SECs were incubated with exogenous H_2_O_2_ (25 μM) for 24 h or Baco II (10 μM) for 72 h. Other SECs were transfected with siNeg or siAQP1. At day 3, all cells were then tested for senescence hallmarks. Left, Upper panel, Baco II or siAQP1 significantly decrease the numbers of SA-β-gal^+^ cells compared to that in siNeg-transfected SECs. Right, quantitative plots are shown for SA-β-gal^+^ cells (%) (n=6). Left, Lower panel, γ-H2A.X immunostaining represent a markedly suppressed DDR in Baco II-and siAQP1-exposed SECs as opposed to that in siNeg-transfected SECs (n=6). **G,** Baco II or siAQP1 restored cell cycle, represented by reduced p16^INK4a^, p19^INK4d^, and p21^WAF1/Cip1^ transcripts in SECs. **H,** A significant decrease in transcription of SASP components IL1α, IL-1β, and IL-6 was seen in Baco II-or siAQP1-exposed SECs compared to siNeg-transfected SECs (n=6). **I,** VCAM1 immunostaining represents its marked downregulation in response to Baco II or siAQP1 in SECs as opposed to that in siNeg-transfected SECs (n=6). Data from *in vitro* cellular experiments represent triplicated biologically independent experiments. Scale bar, 50 and 200 μm. Error bars represent SD (**B-D, F-H**). *P* values were calculated using one-way ANOVA followed by Tukey’s post *hoc* test (**B-D, F-H**). (\**P*<0.05, \*\**P*<0.01, \*\*\**P*<0.001, \*\*\*\**P*<0.0001). Source data are provided as a Source Data file.

By contrast, in replicative SECs, our data revealed that genetic or pharmacological disruption of AQP1 markedly rescues cellular senescence, representing by decreased numbers of SA-β-gal^+^ cells (Fig. 2F) and senescence-related morphological transformations. In line with this, siAQP1 or Baco II strongly preserved telomere length by decreasing γ-H2A.X phosphorylation (Fig. 2F). Quantitative real-time PCR indicated that AQP1 disruption effectively reduces mRNA levels of cell-cycle arrest markers *p16^INK4a^*, *p19^INK4d^*, and *p21^WAF1/Cip1^* (Fig. 2G), and suppressed the SASP profile including *Il1*α, *Il1*β, and *Il6* (Fig. 2D) triggered by replicative senescence. The latter was also confirmed by markedly lower VCAM1 immunofluorescence in SECs exposed to either siAQP1 or Baco II as compared to that in siNeg-transfected SECs (Fig. 2I). In addition, it is confirmed that the AQP1-deficient SECs demonstrate a markedly different Raman spectral peaks and potentially intracellular biochemical content compared to those in siNeg-SECs (Supplemental Fig. 3A,B). Our LDA analysis also affirms distinct biochemical signature of the data point clouds, identifying a low global similarity between the endothelial cell populations (Supplemental Fig. 3C). These spectroscopic dissimilarities hint the potential distinctions in cellular function, metabolic activities, and overall biological heterogeneity within the senescent cell populations. However, the LDA score plot reveals a marked overlap of data point clouds corresponding to siAQP1-SEC and siNeg-PEC populations (Supplemental Fig. 3C).

Together, these data support a potential role of AQP1 in the regulation of EC senescence: AQP1 is essential for maintaining EC proliferating state and may protect these cells against senescence process, whereas AQP1 disruption is suggested to potentially rescue EC senescence.

### AQP1 Differentially Orchestrates Endothelial Energy Supply and Angiogenesis

It has been postulated that endothelial cells undergo permanent alterations in their metabolic and redox states during aging [2,6]. The stable ROS, H_2_O_2_, at higher concentrations, alters mitochondrial dynamics, impairs respiratory chain activity, and disrupts redox homeostasis [2]. It has become clear that disrupted mitochondrial respiration and energy metabolism may drive cellular senescence and dysfunction in these cells [25]. Given the evidence for facilitated H_2_O_2_ transport through AQP1 provided by our (Fig. 1F,G) and others studies, we examined mitochondrial respiration in wild-type and AQP1-deficient PECs and SECs using our standard Seahorse mitochondrial stress test in a high glucose medium. Our assays displayed a significant decrease in basal, and more strikingly maximal, respiration rates (up to 2-fold) in siAQP1-transfected PECs compared with siNeg-transfected PECs (Fig. 3A). Moreover, AQP1 knockdown caused a marked decrease in spare respiratory capacity, with FCCP-stimulated oxidative phosphorylation (OXPHOS) uncoupling, accompanied by reduced ATP biosynthesis (∼1.90-fold) in PECs (Fig. 3A). In contrast to what seen in PECs in the presence of siAQP1 or Baco II, our findings exhibited regulatory role of AQP1 in mitochondrial respiration in an opposite fashion. Accordingly, AQP1 deficiency by siAQP1 or Baco II significantly increased maximal and spare respiration rates (91.4% and 65.5%, respectively) in SECs as opposed to those in siNeg-SECs (Fig. 3B). Furthermore, AQP1-KD could significantly generate higher ATP levels in SECs, which supplies energy for cell migration and angiogenesis in senescent ECs (Fig. 3B). Mitochondrial ATP biosynthesis serves as a critical energy source driving EC migration and angiogenesis by fueling cytoskeletal dynamics and supporting cellular processes essential for vascular network formation [26,27]. In senescent ECs, compromised mitochondrial respiration and reduced ATP production may impair endothelial cell migration and angiogenic capacity, contributing to dysfunctional vascular remodeling [28,29]. Given the differential role of AQP1 orchestrating mitochondrial energy supply, we assessed how AQP1 regulates EC migration and angiogenesis in young and old aortas.

**Fig. 3.**
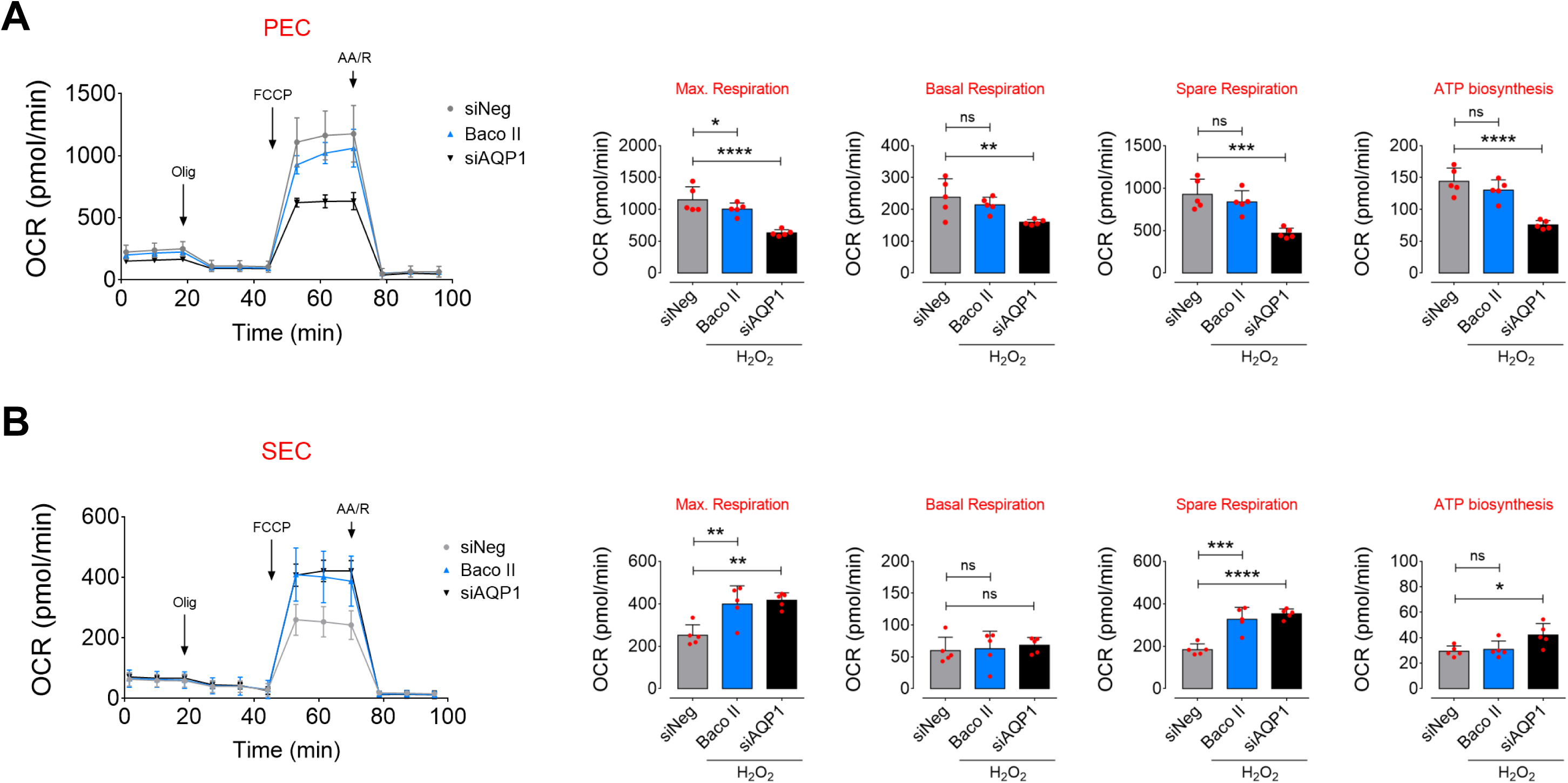
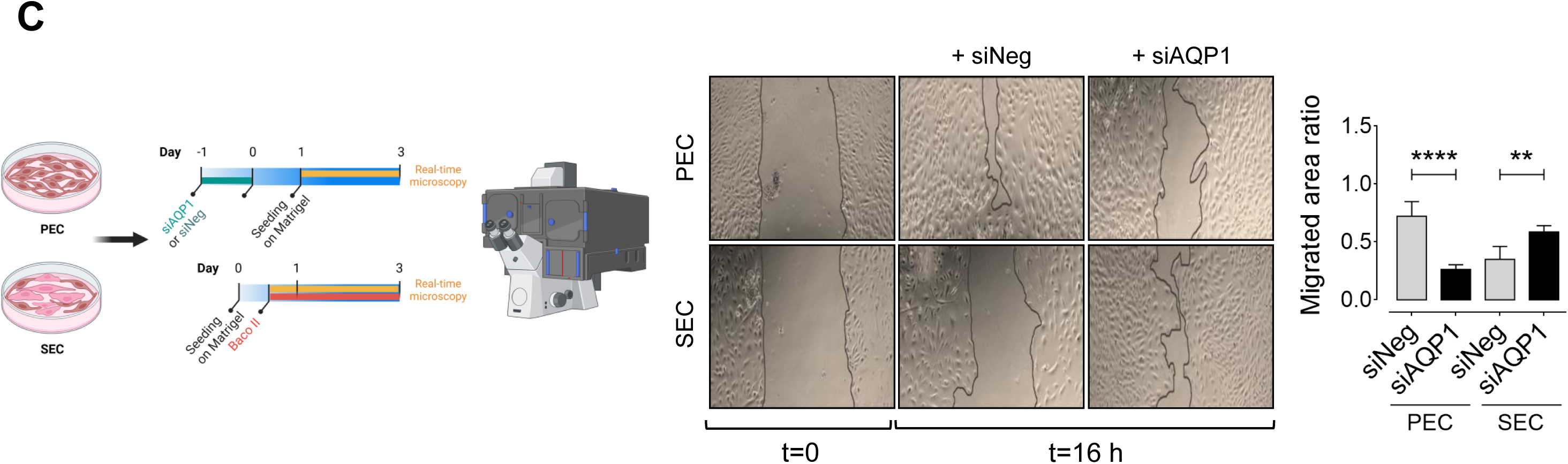
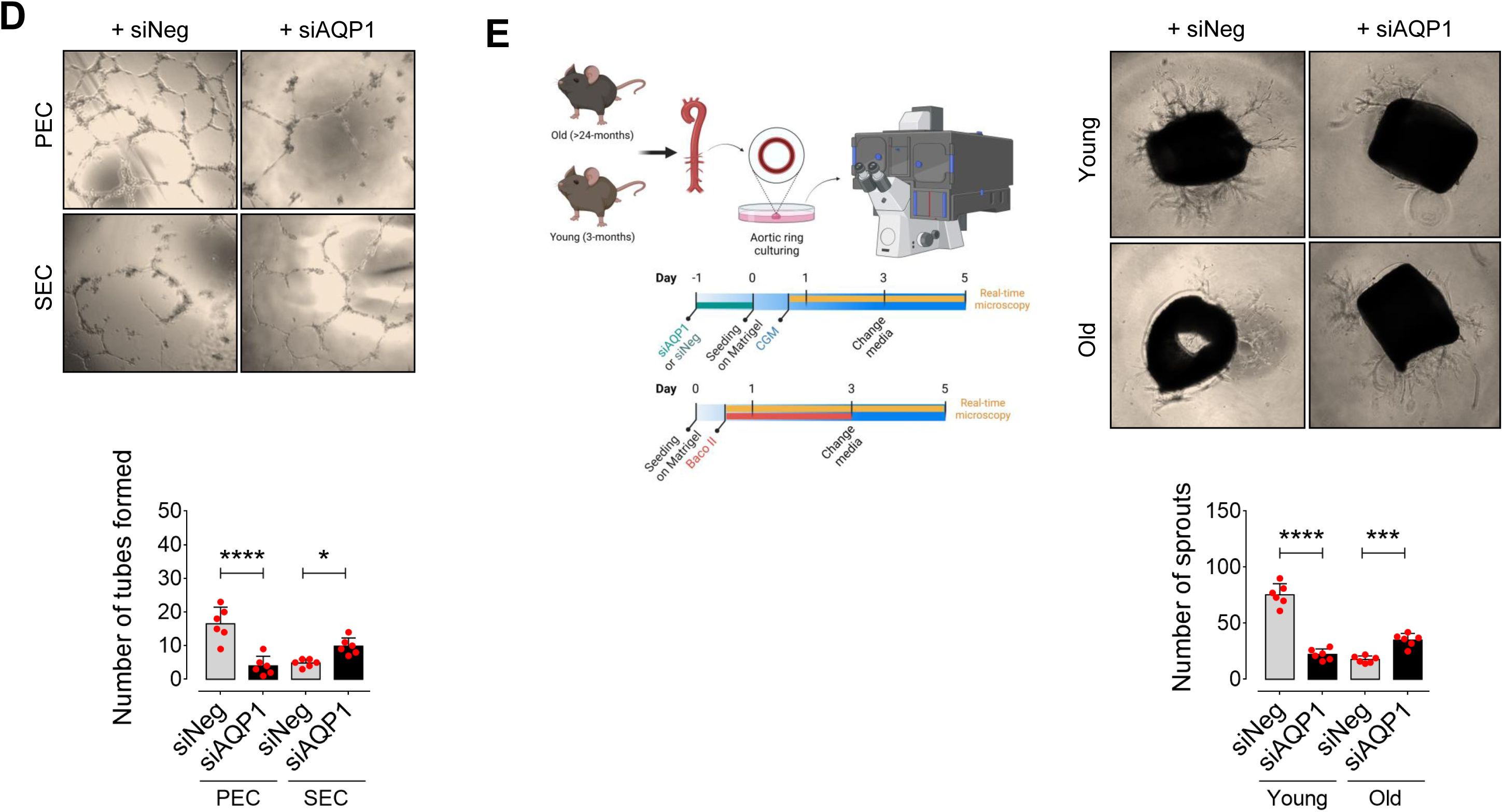
AQP1 differentially orchestrates endothelial energy homeostasis and angiogenesis. Mitochondrial respiratory rates in PECs (**A**) and SECs (**B**) exposed to siNeg, Baco II, or siAQP1 was measured using Seahorse flux analyzer by Cell Mito Stress kit (n=6). Bar charts reveal significant reduction in oxygen consumption rate (OCR) represented by decreased maximal, basal, and spare reserve, and ATP biosynthesis in Baco II-or siAQP1-exposed PECs compared to PECs transfected with siNeg. Conversely, Baco II and siAQP1 restored mitochondrial respiration and markedly increased the OCR parameters in SECs as opposed to those in siNeg-transfected SECs (n=6). **C,** Confocal micrographs depict that siAQP1 significantly reduces cell migration in PECs, shown as larger areas of uncovered surface, to the comparable level seen in siNeg-transfected SECs. While, SECs transfected with siAQP1 exhibited a marked increase in cell migration compared to siNeg-SECs. Right, Bar chart represents the ratio of cell migrated area (n=6). **D,** Confocal images represent the 2-D matrigel tube formation of PECs (Upper) and SECs (Lower) transfected with siNeg or siAQP1. Right, quantitative plot shows a significantly lower number of tubes formed by siAQP1-transfected PECs, while a markedly higher number of tubes was observed in siAQP1-SEC group compared to siNeg-transfected cells (n=6). **E,** Confocal micrographs of the aortic rings from old and young mice show that siAQP1 transfection of young aortas significantly decreases endothelial sprouting, whereas markedly increases angiogenic capacity in old aortas compared to siNeg-transfected aortas. Bottom, quantitative plot is shown for the number of aortic endothelial sprouts (n=6). Scale bars, 200 μm. Data were determined in 6 micrographs from 3 different plates and represent triplicated biologically independent experiments. Error bars represent SD (**A-E**). *P* values were calculated using one-way (**A,B**) and two-way (**C-E**) ANOVA followed by Tukey’s post *hoc* test. (\**P*<0.05, \*\**P*<0.01, \*\*\**P*<0.001, \*\*\*\**P*<0.0001, ns, not significant). Source data are provided as a Source Data file.

Our findings demonstrate a markedly lower migration of siAQP1-transfected PECs (∼2.75-fold), as seen in siNeg-SECs, compared to siNeg-PECs, 16 h post-scratch. However, AQP1 silencing in SECs surprisingly restored cell migration, represented by migrated area ratio, which was greatly impaired during replicative senescence in ECs (Fig. 3C). Next, we examined the effects of AQP1 on tube formation capacity, an *in vitro* measure of angiogenesis, of PECs. In line with prior findings of Saadoun et al. [16], PECs exhibited a significant decrease in the number of tubes formed (∼4-fold) in AQP1-deficient cultures at 16 h. However, siAQP1 could promote tube formation of SECs (103.5%), which was abolished in siNeg-SECs (Fig. 3D). To better define the mechanism of AQP1 for angiogenesis, we established *ex vivo* culture studies using aortas obtained from young and old mice. We demonstrated that young aortic rings, which were transfected with siAQP1 for 48 h, exhibited markedly reduced endothelial sprouting to the magnitude seen in siNeg old aortic rings. On the other hand, AQP1-deficiency greatly restored angiogenic capacity of old aortas, observed as increased total numbers of sprouted endothelial cells, as opposed to those in siNeg old aortic rings at 16 h (Fig. 3E).

### AQP1-regulated EC Senescence is mediated by HDAC4-Mef2A pathway

EC senescence is intricately characterized by profound epigenetic alterations, particularly histone modifications, contributing to dynamic comprehensive changes of gene expression profiles that orchestrate the progression of vascular aging [22,30]. HDAC4 is a key regulator of histone modifications and subsequent expression of the genes involved in senescence-associated phenotypes in endothelial cells [22,31]. Our and others studies suggest that post-translational modification of HDAC4 may modulate histone acetylation states, impacting gene expression patterns associated with endothelial cell senescence and impaired angiogenesis [22,32]. In the present study, our findings exhibited dual role for AQP1 in modulation of HDAC4-regulated signaling pathway in ECs at proliferating and senescence states, through controlling exogenous H_2_O_2_ transport. We observed that AQP1 is crucial for H_2_O_2_ supply, which may regulate HDAC4 post-translational modification and epigenetic alterations, rendering physiological function in PECs. Accordingly, our results showed a marked increase in phosphorylation of CaMKII at Thr286 and its downstream AMPK at Thr172, followed by HDAC4 phosphorylation at Ser632 in PECs exposed to siNeg or H_2_O_2_ (25 μM) (Fig. 4A-D). Using siRNA directed against CaMKII and AMPK, protein expression was suppressed by ∼70-80% compared with negative siRNA-transfected cells for up to 3 days post-transfection, leading to a demonstrable reduction in phosphorylation of AMPK^T172^ and HDAC4^S632^ in PECs, respectively (Supplemental Fig. 4A-C). In siNeg-PECs, physiological H_2_O_2_-mediated HDAC4 phosphorylation facilitated its nuclear export towards the cell cytosol, hence reduces the abundance of HDAC in the cell nucleus as compared to that in AQP1-deficient PECs (Upper panel, Fig. 4G). As a proof of the concept, our phospho-HDAC4^S632^ immunostaining verifies that siAQP1 elicits nuclear localization of HDAC4 in PECs, whereas siNeg-PECS exhibit markedly higher p-HDAC4^S632^ immunofluorescence in the cell cytosol (Fig. 4H). However, senescence-associated decrease in CaMKII-AMPK phosphorylation and the subsequent disruption of HDAC4 phosphorylation at Ser632 in SECs was restored by siAQP1, leading to cytosolic localization of HDAC4 (Lower panel, Fig. 4G) and higher p-HDAC4^S632^ immunofluorescence in the cell cytosol (Fig. 4H). siAQP1-mediated reduction in HDAC4 phosphorylation in PECs has been accompanied by a de-repression of myocyte enhancer factor 2A (*Mef2a*) and subsequent increased Mef2A expression (Fig. 4A,E), which drives an anti-inflammatory phenotype and physiological angiogenic activity of endothelial cells [33]. Our supplementary data supports the notion that HDAC4 is the upstream regulator of Mef2A. As seen in Supplemental Fig. 4D, HDAC4 silencing results in downregulation of Mef2A, consequently reduces eNOS phosphorylation at Ser1177 in PECs. Corroborating our findings, previous studies reveal that HDAC4 phosphorylation on serine residues disrupts HDAC4-Mef2A complex and enhances Mef2A activity, exacerbating tube formation capacity in endothelial cells [32]. Moreover, Mef2A has been recognized for its ability to delay endothelial senescence and, consequently, is considered a potential plasma biomarker for predicting coronary artery diseases in the elderly [34]. Our signaling studies demonstrate that stability of HDAC4-Mef2A complex in AQP1-deficient PECs significantly suppresses eNOS^Ser1177^ phosphorylation (Fig. 4A,F), which is normally seen in replicative senescent ECs. On the other hand, we observed a reverse signaling pattern in SECs. Baco II-or siAQP1-mediated disruption of HDAC4-Mef2A complex could increase Mef2A-regulated eNOS^S1177^ phosphorylation (Fig. 4A,F), which is crucial for physiological EC function and maintaining these cells in proliferating state [22,35].

**Fig. 4.**
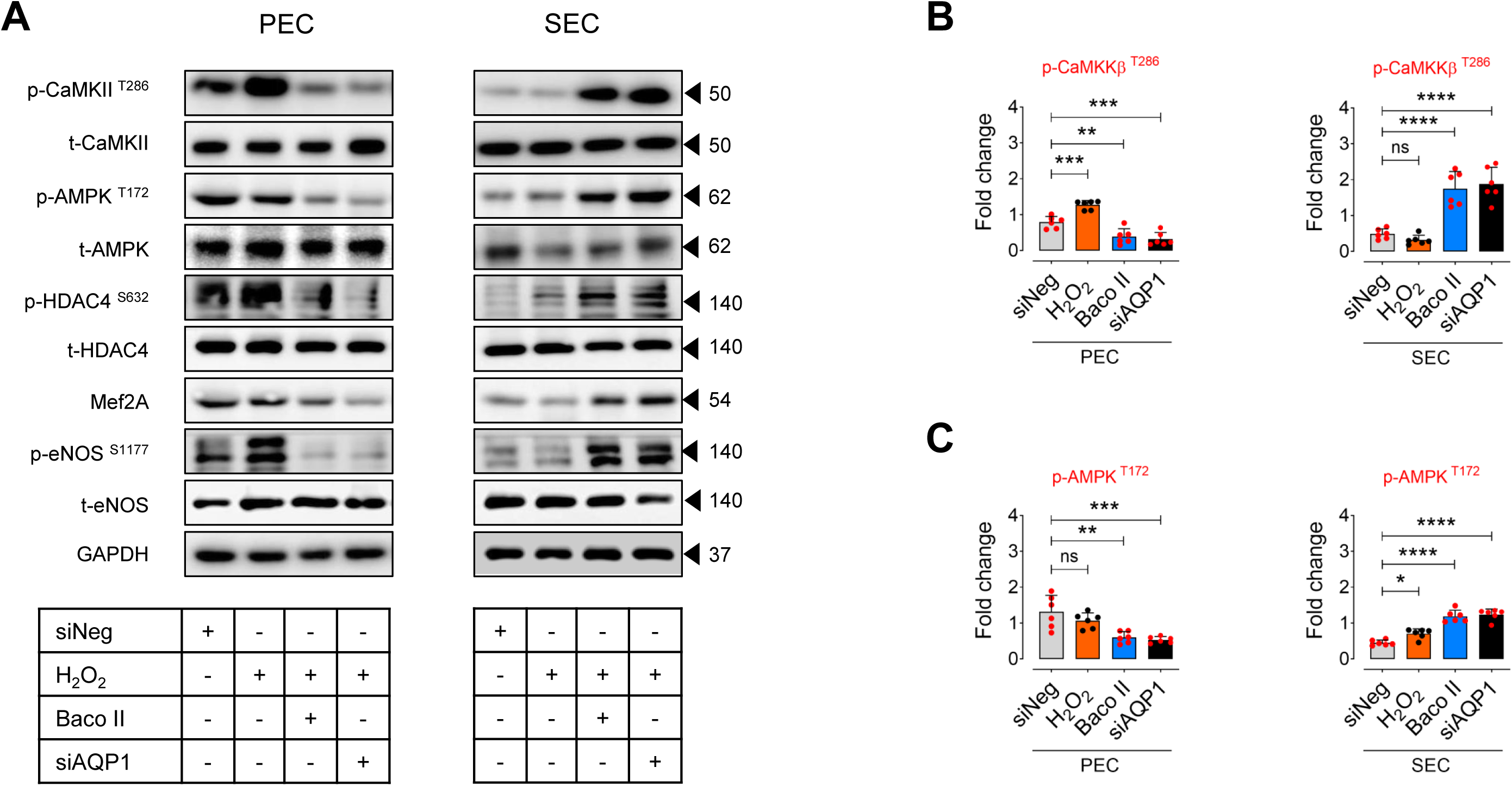
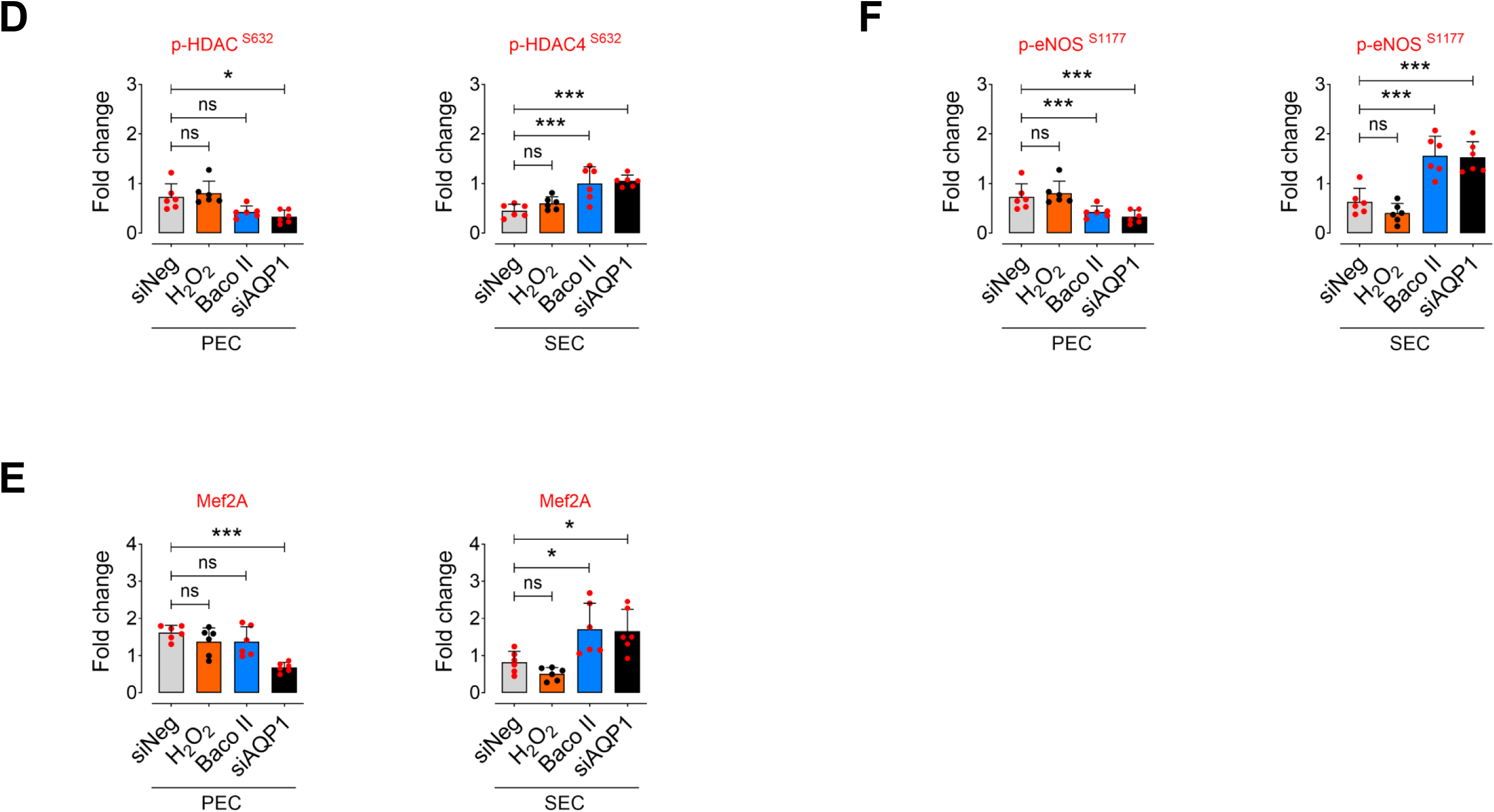
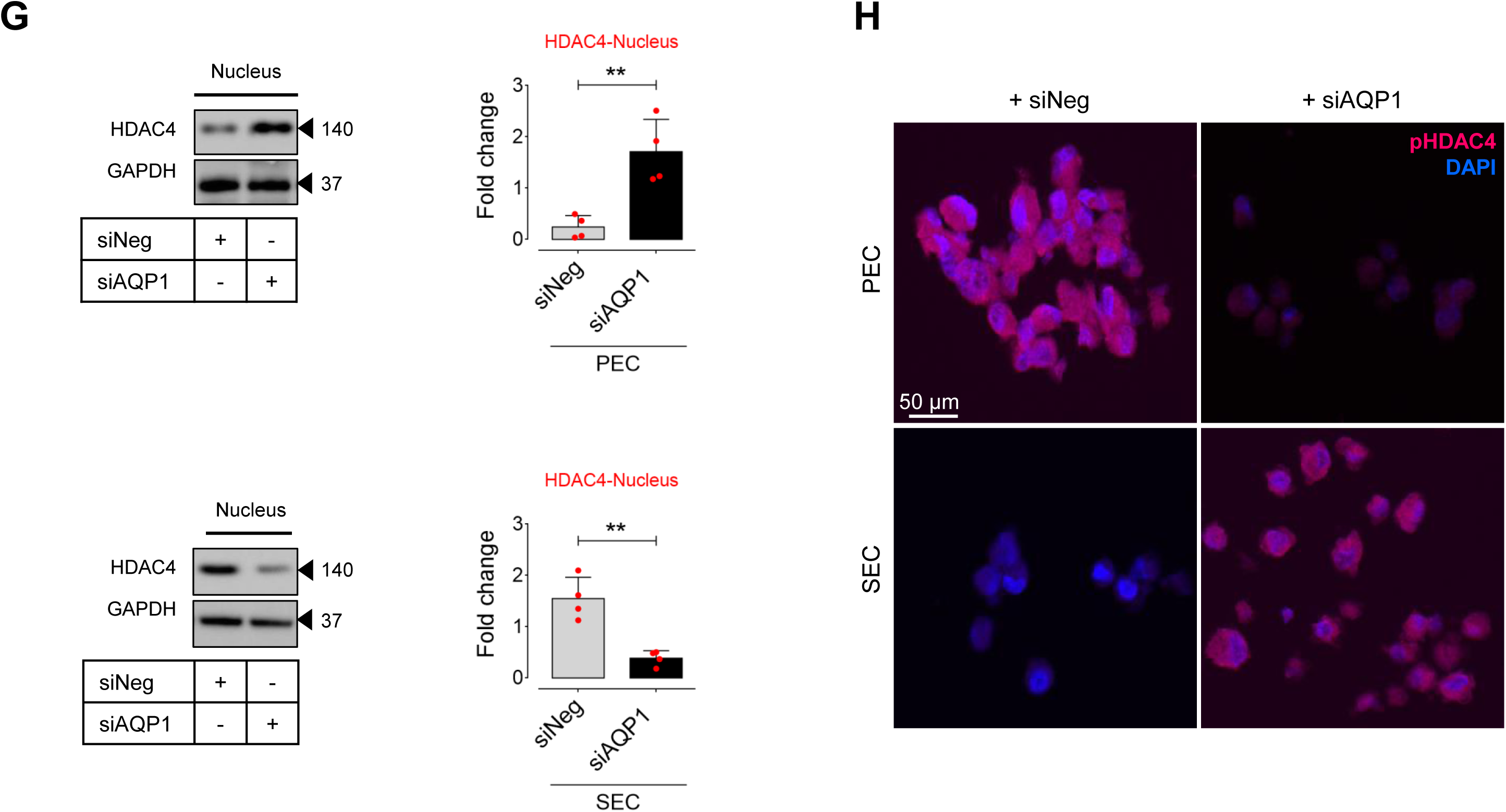
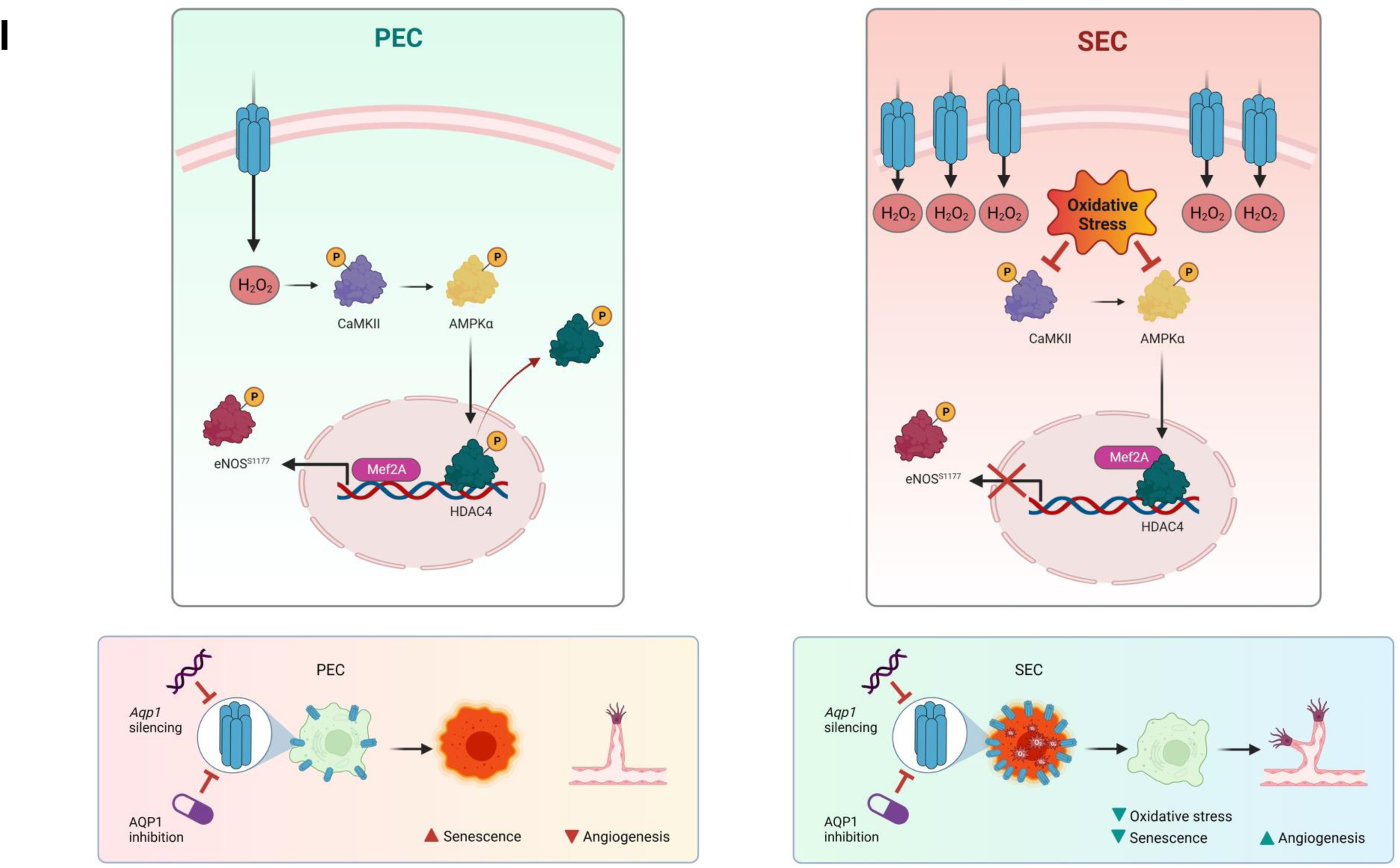
Differential AQP1-mediated endothelial senescence is elicited by HDAC4-Mef2A pathway. **A-F**, Immunoblot analysis reveals the differential role of AQP1 in the regulation of CaMKII-AMPK-mediated phosphorylation of HDAC4 at Ser632 and the subsequent Mef2A-eNOS phosphorylation in PECs *versus* SECs (n=6). **G,** Immunoblots (left) and quantitative plots (right) reveal that siAQP1 stimulates nuclear localization of HDAC4, represented as increased expression in the nucleus, in PECs (top), while it facilitates nuclear export, represented as a decrease in HDAC4 expression in the cytosol, in SECs (bottom) (n=6). **H,** p-HDAC4 immunostaining shows that siAQP1 markedly decreases HDAC4 phosphorylation, facilitating its nuclear localization in PECs. However, siAQP1 transfection facilitates HDAC4 translocation towards the cell cytosol, shown as higher pHDAC4 fluorescence, in SECs (n=6). **I,** The mechanism underlying the differential contribution of AQP1 to endothelial cell senescence and angiogenic capacity. AQP1 differentially orchestrates H_2_O_2_-mediated EC senescence. (Left) In PECs, AQP1 is crucial for H_2_O_2_-regulated CaMKII-AMPK stimulation and the subsequent HDAC4 phosphorylation (nuclear export), facilitating Mef2A expression and eNOS^S1177^ phosphorylation. *Aqp1* silencing or pharmacological inhibition of AQP1 promotes cellular senescence and impairs angiogenesis in these cells. (Right) In SECs, higher AQP1 abundance contributes to oxidative stress that suppresses CaMKII-AMPK-mediated HDAC4 translocation, subsequently leading to reduced Mef2A expression and eNOS activation. Conversely, AQP1 deficiency reduces intracellular ROS accumulation and thereby both reverses cellular senescence and restores angiogenic capacity in these cells. Scale bars, 50 μm. Data were determined in 6 micrographs from 3 different plates and represent triplicated biologically independent experiments. Error bars represent SD (**B-G**). Continuous data are presented as mean ± SD. Statistical analysis was performed with a two-tailed unpaired Student’s *t*-test (**H**) and one-way ANOVA followed by Tukey’s post *hoc* test (**B-F**). (\**P*<0.05, \*\**P*<0.01, \*\*\**P*<0.001, ns, not significant). Source data are provided as a Source Data file.

Our data support the notion that AQP1 differentially regulates protein kinases-mediated subcellular localization of HDAC4 and its control on Mef2A-nitric oxide (NO) signaling axis in proliferating and senescent ECs.

## Discussion

In the current study, we found that aging enhances the distribution of aquaporin 1 in aortic endothelial cells in mice. It was also verified in replicative senescent ECs that exhibit higher AQP1 expression. Perhaps the most striking finding is that AQP1 differentially orchestrates extracellular H_2_O_2_-induced cellular senescence in ECs by upregulating CDK inhibitors, SASP, and DNA damage responses. We also suggested a new differential role for AQP1 in driving intrinsic angiogenic incompetence in replicative SECs that is integral to vascular dysfunction. Our combined cellular senescence, metabolic, and angiogenesis studies revealed that AQP1 is crucial for a persistent proliferation and angiogenic capacity in ECs. Accordingly, *Aqp1* knockdown or selective blockade of AQP1 induces morphological and mitochondrial alterations in aortic ECs that lead to cellular senescence and consequently impairment of angiogenesis. By contrast, age-related increase in AQP1 expression promotes senescence phenotype and impaired energy homeostasis, thereby fostering angiogenic incompetence. Our signal transduction studies revealed that AQP1-mediated H_2_O_2_ transport differentially modulates post-translational modification of multiple protein kinases regulating HDAC4, as an epigenetic regulator of endothelial function and angiogenesis. The HDAC4 (de)phosphorylation shuttles between the cell cytosol and nucleus, which thereby regulates Mef2A transcription and subsequent eNOS biology. Corroborating these results, our and others’ previous studies demonstrated that accumulation of senescent ECs plays a causative role in pathogenesis of vascular diseases such as atherosclerosis [6,36]. Therefore, our findings provide a key pathophysiological mechanism for the induction of endothelial senescence and the development of atherosclerosis upon aging. Moreover, genetic or pharmacological AQP1 disruption can be potentially suggested as a potential anti-senescence strategy for improvement of atherosclerosis during aging.

Aquaporins have long been recognized as water transporters governing osmotic water movement in mammalian cells [37]. Among the 13 known AQP isoforms, AQP1, AQP3, AQP8, and AQP9 have been identified to potentially transport H_2_O_2_, expanding their role beyond water transport through their intrasubunit pores [38]. Structural analyses in both humans and mice support AQP1’s dual capacity for transporting H_2_O_2_ and water [14]. While H_2_O_2_ was traditionally believed to freely diffuse across the plasma membrane, our HyPer7.2 real-time imaging in endothelial cells, along with prior studies in cardiac myocytes [14], suggests a specific role for AQP1 in directing H_2_O_2_ transport to various subcellular compartments. AQP4, alongside AQP1, plays a role in osmotic water transport within these cells [39]. Consequently, the deletion of *Aqp1* may not significantly impact water movements. Notably, the fact that AQP4 lacks the ability to transport H_2_O_2_ both elucidates the distinct role of AQP1 in H_2_O_2_-regulated cellular events and implies that the transport of H_2_O_2_, rather than water, by AQP1 is pivotal for modulating cell signaling. Although little is known about the impact of aging on AQP1 expression in vascular cells, studies showed increased AQP1 expression in cardiac myocytes during aging, contributing to myocardial ischemia in the elderly [19]. Our current findings reveal that aging is associated with higher expression of AQP1 in aortic ECs in mice exhibiting EC dysfunction. Our and others’ prior studies demonstrated that H_2_O_2_ drives epigenetic alterations and SASP, triggering EC senescence and dysfunction [8,21,22]. Given that the accumulation of senescent ECs underpins the pathogenesis of atherosclerosis and atherosclerotic cardiovascular diseases [36], AQP1’s significance extends beyond its impact on osmotic water permeability in the regulation of EC biology in aging.

Our findings show that, under proliferating conditions, the expression of AQP1 is relatively low in ECs whereas it increases in replicative senescence ECs. This apparently boosts the transport of H_2_O_2_ across membrane boundaries, thereby accumulating H_2_O_2_ within the SECs. H_2_O_2_, at physiological levels, stimulates EC proliferation and migration as well as tube formation in an in vitro model of angiogenesis [40]. It has been postulated that low concentrations of H_2_O_2_ are involved in numerous signal transduction pathways, including CaMKII-eNOS, fostering angiogenic capacity [41]. Corroborating with this, our results demonstrate that *Aqp1* knockdown or AQP1 blockade inhibits H_2_O_2_ transport into PECs and remarkably reduces *in vitro* cell migration and tube formation as well as *ex vivo* endothelial sprouting in young aortas. It highlights the pivotal role of AQP1 in supplying physiological H_2_O_2_ from extracellular non-enzymatic and enzymatic sources, such as NOX2 and xanthine oxidase [42], for regulating EC migration and angiogenesis. In addition, the AQP1-deficient PECs exhibited cellular senescence hallmarks encompassing increased SA-β-galactosidase activity and proliferative arrest, represented by increased expression of the CDK inhibitors p16^INK4a^, p19^INK4d^, and p21^WAF1/Cip1^, along with DNA damage response (increased γ-H2A.X phosphorylation). Our *in vitro* studies strongly demonstrate excess SASP responses by upregulating IL1α, IL1β, IL6, and VCAM1 in these cells. These findings are consisted with the literature that indicates the biphasic role of H_2_O_2_ in regulating EC functions. The role of hydrogen peroxide (H_2_O_2_) in cellular senescence is complex and context-dependent. H_2_O_2_, at physiological concentrations, induces oxidative eustress, while transient or sustained increases in H_2_O_2_ levels may dysregulate redox homeostasis, leading to chronic oxidative distress [43,44]. Therefore, ensuring the preservation of physiological H_2_O_2_ levels is essential for sustaining cellular processes that maintain ECs in a proliferative state.

During aging, the accumulation of excessive ROS in the extracellular space enhances the levels of advanced oxidation protein products (AOPP) in ECs, which have been identified to accelerate senescence process [45]. For instance, in atherosclerosis-prone apolipoprotein E-deficient (*ApoE^−/−^*) mice fed a high-fat diet, AOPP attenuated autophagy in aortic ECs, which leads to p53 SUMOylation at K386, ultimately resulting in premature senescence [46]. H_2_O_2_-induced oxidative stress, along with suppressed activity of antioxidant defense, has been found to stimulate p53-mediated mitochondrial oxidative stress through destroying mitochondrial membrane potential and dysregulated energy metabolism, triggering accelerated senescence and dysfunction, proven as subclinical stage of atherosclerosis [47]. Corroborating our findings, excessive H_2_O_2_-induced oxidative stress shuts down mitochondrial oxidative phosphorylation, resulting in impaired ATP biosynthesis in SECs [22, 48]. This substantiates the occurrence of mitochondrial metabolic alterations in these cells, potentially contributing to diminished cell migration and compromised angiogenic capacity. Not surprisingly, the heightened SASP observed in SECs, characterized by increased levels of IL1α, IL1β, IL6, and VCAM, may be attributed to ROS-induced mitochondrial damage and widespread mitochondrial outer membrane permeabilization (MOMP). This process facilitates the release of mitochondrial DNA (mtDNA) into the cell cytosol. The presence of mtDNA in the cytosol activates the cGAS–STING pathway, thereby orchestrating SASP components such as IL6 and IL8, ultimately promoting inflammatory responses [49]. Therefore, the inhibition of H_2_O_2_ transport into ECs can suggest a potential strategy to prevent EC senescence and dysfunction. Accordingly, we showed that both Bacopaside II-mediated AQP1 blockade and *Aqp1* knockdown ameliorates H_2_O_2_-induced oxidative stress and the subsequent senescence hallmarks in SECs. In the AQP1-deficient SECs, we observed a restored proliferative arrest capacity, along with marked reduction in cell-cycle arrest markers including p16^INK4a^, p19^INK4d^, and p21^WAF1/Cip1^, and DNA damage, which is probably mediated by downregulating p16^INK4a^ and p19^INK4d^. Consistently, p16^INK4a^-expressing ECs demonstrate DNA damage features, whereas elimination of the senescent ECs delays the incidence of atherosclerosis and subsequently extends the median lifespan in progeroid mice [50]. Our findings further demonstrated that AQP1 deficiency restores the profound reductions in maximal and spare mitochondrial respiration seen in the SECs, suggesting that it reinforces these cells to compensate for decreased ATP biosynthesis to maintain cellular energy homeostasis essential for angiogenesis. In line with this, restored mitochondrial respiration and energy supply in Bacopaside II-or siAQP1-treated SECs is accompanied by a high energetic proliferating state, governing a marked increase in pro-angiogenic efficiency. Additionally, our AQP1-deficiency interventions restored aortic endothelial sprouting *ex vivo* in old mice, supporting cytoskeletal dynamics and formation of new blood vessels for recovery of ischemic vascular diseases in older adults [51]. In these clinical scenarios, impaired angiogenesis fails to elicit the necessary compensatory mechanisms required for augmenting blood supply in conditions characterized by reduced blood flow and compromised tissue perfusion [52]. This deficiency underscores the urgency of understanding and harnessing angiogenesis as a successful treatment of aging-related cardiovascular pathologies.

Here, we grappled with the endeavor to unravel a fundamental question: “how does AQP1 differentially modulate cellular function in both proliferating and senescent ECs?” Based on the literature, AQP1 is co-localized with enzymatic sources of ROS, including NOX2, in caveolae-enriched membrane fractions that enhance ROS-sensitive kinases signaling cascades [14]. These kinases, in a downstream manner, are instrumental in eliciting cellular senescence and dysfunction processes. Hence, our inquiry focused on elucidating the answer to this question through an examination of H_2_O_2_-regulated kinases that modulate eNOS signaling in ECs. We first observed that protein kinase CaMKII is phosphorylated at Thr286, which thereby stimulates the phosphorylation of its downstream energy sensing and signaling regulator AMPK at Thr172 in the presence of physiological levels of H_2_O_2_ in PECs. This was followed by post-translational modification of HDAC4, the epigenetic regulator of eNOS signaling pathway. Our findings showed that AQP1 is essential for HDAC4 phosphorylation at Serine site 632, which subsequently controls its nuclear export, in proliferating ECs. It returns to the fact that redox/metabolic status in ECs orchestrates permanent changes in their epigenetic programming [2]. The CaMKII-mediated hierarchical phosphorylation of HDAC4^S632^ facilitates its subcellular localization [22]. In accordance to our and others’ previous studies, H_2_O_2_ regulates HDAC4 shuttle between cell cytosol and nucleus through two potential mechanisms, one direct oxidation of HDAC4, and another one H_2_O_2_-regulated phosphorylation of HDAC4 in ECs [22,32]. Matsushima and colleagues discovered that phosphorylation of HDAC4, secondary to its oxidation, facilitates nuclear export in cardiac myocytes [53]. Moreover, oxidized HDAC4 has been shown to interact more easily with protein kinases of the CaMKII family. Consistently, our prior investigation demonstrated that gut-derived phenylacetic acid, by NOX4-mediated H_2_O_2_ generation, both oxidizes and phosphorylates (through CaMKII stimulation) HDAC4, facilitating its cytosolic accumulation in PECs [22]. We next observed that HDAC4 nuclear export facilitates de-repression of Mef2A in PECs. It has been previously postulated that HDAC4 physically interacts with Mef2A and forms HDAC4-Mef2A complex, which is potentially disrupted by H_2_O_2_ in ECs [32]. Indeed, unbinding of HDAC4 to Mef2 in the nucleus causes an increase in *Mef2A* transcriptional activity, leading to an anti-inflammatory state and increased tube formation potential in the cells [32]. In PECs, de-repression of Mef2A was accompanied by increased phosphorylation of eNOS at specific site Ser1177, which has long been recognized to play a crucial role in physiological EC function and angiogenesis. Conversely, in AQP1-deficient PECs, lower intracellular H_2_O_2_ contributes to a marked decrease in phosphorylation of protein kinase CaMKII and its downstream AMPK. Both H_2_O_2_ loss and reduced activation of CaMKII-AMPK inhibit the HDAC4 post-translational modification and the subsequent disruption of HDAC4-Mef2A complex in the nucleus, ultimately suppressing phosphorylation responses on eNOS^Ser1177^. As reported previously, genetic deletion of eNOS disrupts VEGF-induced angiogenesis *in vivo*, whereas phosphorylation of eNOS at Ser1177, but not Thr495, by multiple protein kinases increases neovascularization in ischemic tissues [54]. The findings, therefore, boost our understanding of the primordial role of AQP1, as a redox-epigenetic regulator of endothelial function in a youthful state. However, it elucidates only a sequence of the whole scenario of AQP1 in the regulation of EC senescence and function. Our findings reveal that AQP1 plays a differential role in orchestrating Mef2A-eNOS signaling in SECs compared to PECs. Specifically, SECs expressing higher levels of AQP1 exhibit a notable decrease in phosphorylation responses at specific sites on CaMKII and AMPK (Thr286 and Thr172, respectively). Excessive H_2_O_2_ and other ROS within SECs disrupt the phosphorylation responses of these protein kinases. This observation aligns with our recent studies indicating that CaMKII phosphorylation declined in SECs, in which mitochondrial ROS induces oxidative stress [22]. The diminished CaMKII phosphorylation due to oxidative stress can impair its downstream signaling cascades, contributing to endothelial dysfunction [55]. Accordingly, prior *in vitro* studies revealed that higher concentrations of H_2_O_2_ (>10 μM) inhibit AMPK phosphorylation at Thr172 across various human endothelial cell types [56]. Additionally, exposure to H_2_O_2_ induces morphological changes indicative of cellular senescence in NIH-3T3 cells within three to five days, characterized by enlarged and flattened morphology. This senescent phenotype is accompanied by a notable decrease in phosphorylated AMPKα (Thr172) levels and decreased activity, alongside reduced transcriptional levels of genes regulating fatty acid β-oxidation. Conversely, activation of AMPK has shown potential in preventing H_2_O_2_-induced senescence in these cells [57]. In the presence of excess H_2_O_2_-induced oxidative stress and subsequent absence of CaMKII-AMPK activation, HDAC4 becomes dephosphorylated and thereby accumulated in the nucleus at the magnitude seen in AQP1-deficient PECs. Indeed, HDAC4 dephosphorylation is a prerequisite for Mef2A repression and further decrease in eNOS^Ser1177^ phosphorylation in SECs that exhibited a disrupted angiogenesis. Our findings, in turn, revealed that AQP1 blockade or *Aqp1* knockdown, by upregulating CaMKII-AMPK phosphorylation, enhances HDAC4^S632^ phosphorylation and stabilizes its cytosolic localization, which thereby increases Mef2A expression in SECs. Indeed, the outcome was an increase in eNOS phosphorylation at Ser1177, which can regulate endothelial sprouting and neovascularization *in vivo* through enrichment of genes related to cell polarity such as Partitioning Defective (PARD) family detected at the single-cell level [58]. eNOS has also been shown to regulate angiogenesis in a hind limb ischemia model, in which the EC-specific knockout of Rac1, a Rho family member, reduces eNOS activity and NO generation that finally abolishes aortic capillary sprouting and peripheral neovascularization [59]. Therefore, in addition to restoration of mitochondrial respiration and energy supply, AQP1 deficiency may restore angiogenic capacity of ECs likely through disrupting HDAC4-Mef2A complex and increasing eNOS function. Our findings suggest AQP1 blockade or *Aqp1* deletion as effective senotherapeutic strategy with potential of pro-angiogenic efficiency for recovery of ischemic vascular diseases such as coronary artery disease and peripheral arterial disease in older adults.

Taken together, our findings reveal that peroxiporin AQP1 is a key regulator of cellular senescence and angiogenic capacity in endothelial cells through intracellular H_2_O_2_ modulation and subsequent differential regulation of epigenetic signaling and mitochondrial function in aging. We also propose that AQP1 disruption holds a promise to rescue replicative senescence and restore energy homeostasis and function in endothelial cells. Our studies open an avenue to complement the senotherapeutic armamentarium with specific AQP1 blockade for the prevention of endothelial senescence and improvement of angiogenic potential in older adults at higher risk for ischemic vascular diseases.

## Methods

### Mice

Our research complies with all ethical regulations approved by the Institutional Animal Care and Use Committee at University of Zurich. Animal care and all experimental protocols were in accordance to the Directive 2010/63/EU of the European Parliament and of the Council of 22 September 2010 on the protection of animals used for scientific purposes.

8 week old wild-type C57BL/6J female and male mice were purchased from the Jackson Laboratory and maintained in our facilities for specific times for normal aging modeling. All mice were individually housed in controlled environments in plexiglass cages under a strict 12 h:12 h light/dark cycles at an ambient temperature of 23 ± 1°C and humidity of 55 ± 10%, and fed standard chow diet (including 19% Protein, 61% Carbohydrate and 7% Fat, #D11112201, Research Diets, New Brunswick, NJ, USA) until 3 (as Young) or >24 (as Old) months of age. Mice had access to drinking water and food *ad libitum*. The animals were monitored for body weight during their lifespan. The investigators in this study worked in a double-blinded manner.

### Tissue harvesting

Mice were euthanized by CO_2_ inhalation in an area separate from the housing facility. Heart was perfused with normal saline and aorta was harvested, followed by removal of adhering connective and fat tissues. Aortas were kept in Opti-MEM + 2.5% (v/v) fetal bovine serum (FBS) for *ex vivo* vasorelaxation and aortic endothelial sprouting. For immunohistochemical and signaling analyses, aortic tissues were quickly fixed in 10% formalin or snap-frozen in liquid nitrogen followed by storage at-80 LC, respectively.

### Cell Line

The primary human aortic endothelial cells (HAEC) were grown in EBM-2 endothelial cell growth basal medium supplemented with EGM-2 cell growth supplement pack containing FBS (10% v/v), L-glutamine (2 mM) and penicillin–streptomycin (100 μg/mL) and incubated at 37°C in 5% CO_2_. The cell line was tested to exclude any positive status for mycoplasma. Cells were studied in passages 4-5 (proliferating) and 15-17 (replicative senescence). The cellular senescence was verified by SA-β-galactosidase staining and immunoblots of p21 ^WAF1/Cip1^ [22].

### Endothelial Cell Senescence Models

#### Replicative senescence

HAEC (p.4 or p.5) were seeded in 100% EGM-2 at a density of 5,000 cells/cm^2^ and the culture medium was changed every 48 h. To generate the corresponding proliferative control (PEC), cells were trypsinized and cultured in a growth medium for 48 h to proliferate. In parallel, we continued cell passage every 48 h until the proliferation is suppressed and replicative senescence phenotype (due to repetitive passages) is proved (SEC; p.15 to p.17). The cellular senescence was verified by SA-β-galactosidase staining.

#### H_2_O_2_-induced premature senescence

In order to induce premature senescence, proliferating HAEC (p.4 or p.5) were cultured in 100% EGM-2 with 10% fetal bovine serum (FBS) and treated with exogenous H_2_O_2_ (50 μM). After 4 h, the culture medium was changed, and cells were kept on culture for up to 72 h.

### SA-**β**-galactosidase staining

Cultured HAEC cells were stained for SA-β-galactosidase according to the manufacturer’s protocol (Merck Millipore). Briefly, cells were fixed for 15 min and then stained with SA-β-gal detection solution overnight at 37°C. Senescent cells, illuminated as blue-stained cells, were captured under a light Olympus microscope and analyzed using ImageJ software.

### CellRox green staining

Cells were grown to confluence in growth medium and treated with CellROX^®^ Green reagent at a final concentration of 5μM (Invitrogen, C10444) for 30 min at 37 °C to detect intracellular ROS by a Leica TCS-SP8 fluorescence microscope. The fluorescence intensity was evaluated using ImageJ software.

### HyPer7.2 fluorescence imaging

The cells expressing adenovirus serotype 5 (AV5)-HyPer7.2 targeted to the cell cytosol at a MOI of 1000 were treated with siNeg, siAQP1 or Bacopaside II. Cells were then washed and incubated in a HEPES-buffered solution containing 140 mM NaCl, 5 mM KCl, 2LmM CaCl_2_, 1LmM MgCl_2_, 10 mM D-glucose and 1 mM HEPES (pH 7.4) for 2 hours at 37 °C. Coverslips were mounted on a live-cell imaging platform that allowed for stable superfusion.

For real-time fluorescence imaging, the ratiometric HyPer7.2 biosensor was excited at 420 nm and 490 nm, and emission was recorded at 530 nm using Metafluor Software, as previously described [10,35,60]. The cytosolic H_2_O_2_ measurement was performed with a 20X oil immersion objective. Following background subtraction, mitochondrial H_2_O_2_ was defined as R/R_0_, ratio, where R is the ratio of the 490 nm to the 420 nm signals and R_0_ is the baseline 490/420 ratio.

### siRNA transfection

HAECs were grown to 70–80% confluence for transfection with OnTARGETplus short-interfering RNA (siRNA) smart pool (Dharmacon) targeting AQP1 (siAQP1, L-021494-00-0005), AMPKα (siAMPKα, L-005027-00-0005), and Mef2a (siMef2a, L-009362-00-0005). OnTARGETplus Non-targeting scrambled control siRNA (siNeg, D-001810-10-05) was utilized for as negative control. Cells were transfected with siRNA using Lipofectamine RNAiMAX transfection reagent (Invitrogen) according to the manufacturer’s protocols. After 48 h, HAECs were collected for subsequent analysis. The silencing efficiency was confirmed by immunoblots.

### Antibodies

Antibodies against phospho-AMPKα Thr172 (1:1000; 2535), total-AMPKα (1: 1000; 2532), phospho-eNOS Ser1177 (1:1000; 9517), phospho-eNOS Thr495 (1:1000; 9574), total-eNOS (1: 1000; 32027), Histone H3 (1:1000; 9715), Mef2a (1:1000; 9736), and phospho-Histone H2A.X Ser139 (1:100 and 1:1000; 80312) were purchased from Cell Signaling Technology; antibodies against VCAM-1 (1:100, 1:200 and 1:1000; MA5-31965), and VE Cadherin (1:100; PA5-19612) were obtained from Invitrogen; antibody against AQP1 (1:750 and 1:1000; ab168387), phospho-HDAC4 Ser632 (1:250 and 1:1000; ab39408), and total-HDAC4 (1:1000; ab12172) were obtained from abcam; antibody against CD31 (1:50; DIA-310) was purchased from Dianova; antibody against H3ac (Pan-Acetyl) (1:500; sc-518011) was obtained from Santa Cruz Biotechnology; antibody against p16^INK4A^ (1:100; ZRB1437) was purchased from Merck Millipore; antibodies against goat anti-rabbit IgG-HRP (1:2000; 4030-05) and goat anti-mouse IgG-HRP (1:2000; 1036-05) were purchased from Southern Biotechnology.

### Immunoblotting

Proteins were extracted from in ice-cold RIPA lysis buffer (ThermoFisher Scientific) or nuclear and cytoplasmic extraction reagents (for protein isolation from nuclear and cytosolic fractions) supplemented with protease and phosphatase inhibitors cocktails (ThermoFisher Scientific). Equal amounts of protein lysates (20 µg) were separated in 10% SDS-PAGE gels and transferred to PVDF membranes (Bio-Rad). The membranes were incubated with specific primary antibodies (1:1000) overnight at 4°C and then with corresponding horseradish peroxidase (HRP)-labeled secondary antibodies (1:2000) for 1h. The protein bands were visualized using an Immobilon Western HRP substrate Crescendo and Forte Reagents and Amersham Imager 600 (GE Healthcare). Quantitative densitometric analyses were performed using ImageJ software.

### Real-time quantitative PCR

Total RNA from cells and aortas was extracted using TRIzol Reagent^®^ (Sigma-Aldrich), according to the standard protocol previously described [61]. cDNA was synthesized using High-Capacity cDNA Reverse Transcription Kit (Thermo Fisher Scientific) and amplified in a StepOnePlus RT-PCR thermocycler (Applied Biosciences) with Power SYBR Green PCR Master Mix (Thermo Fisher Scientific). Genes of interest were amplified with the corresponding gene-specific pairs of primers designed according to the coding strand of genes in the National Center for Biotechnology Information (NCBI) database using Integrated DNA Technologies (idtdna) and listed in Supplementary Table 1.

### Opal multiplex immunofluorescence

Murine aortas and aortic endothelial cells were formalin-fixed and paraffin-embedded. Experimental sections (4-μm thickness) and positive tissues were stained with specific primary antibodies followed by Opal multiplex immunostaining system (Akoya Biosciences) to generate the stained slides. The staining conditions for all primary antibodies were optimized using chromogenic DAB detection (Leica Biosystems). Using an Opal 7-Color Automation IHC Kit (Akoya Biosciences), Opal singleplex and multiplex for initially-optimized antibodies AQP1, CD31, p16^INK4A^, VCAM1, and γ-H2A.X^Ser139^, as well as DAPI (for nuclear counterstaining) were conducted on a Leica Bond Rx automated autostainer (Leica Biosystems). The panel of primary antibody/fluorophore pairs was applied to spectrally co-localize them in the same cellular compartment (AQP1/Opal 480, CD31/Opal 480, p16^INK4A^/Opal 520, VCAM1/Opal 570, γ-H2A.X^Ser139^/Opal 690), as recommended by the manufacturer. All multiplex immunofluorescence slides were scanned on a Vectra Polaris Automated Quantitative Pathology Imaging System (Akoya Biosciences) at ×20 magnification, which generates a single unmixed image tiles using Phenochart (Version 1.2.0, Akoya Biosciences). Images were automatically stored on a secure networked server for further analysis.

### Seahorse Mito Stress assay

HAECs were seeded at a density of 5,000 cells/well per on Seahorse XF24 tissue culture plates (Seahorse Bioscience). On the day of experiment, cells were washed and medium was replaced with culture medium supplemented with 25 mM glucose, 2 mM glutamine and 1 mM pyruvate (pH 7.4). For a standard mitochondrial stress test, oligomycin (1 μM), an ATP synthase blocker, FCCP (3 μM), oxidative phosphorylation uncoupler, and Antimycin A/Rotemnone (0.5 μM for each), complex III/ complex I inhibitor, are injected to assess oxygen consumption rate (OCR) and extracellular acidification rate (ECAR) over a 3-min period. At any condition, 5 consecutive measurements of OCR and ECAR are done. Data are normalized to total protein/ well.

### Single-Cell Micro-Raman spectroscopy

HAECs (both proliferating and replicative senescent cells) were seeded on sterile sterile μ-channel slide with a 0.17 mm thick borosilicate glass bottom (BBiospex^®^, microphotonX GmbH, Tutzing, Germany) to 70-80% confluence in EGM-2 with 10% FBS for further transfection with siNeg and siAQP1. After 48 h, the cells were washed with PBS and fixed with 4% paraformaldehyde (PFA) for 15 min at 4°C. Cells were subjected to Raman spectroscopy using the Raman microscopic-laser trapping system (Biospex^®^, microphotonX GmbH, Tutzing, Germany) with an inverted microscopic setup. Raman spectra were captured using a laser (at 785 nm and 80 mW laser power; TOPTICA Photonics AG, Graefelfing, Germany) for 10-sec acquisition, with 3 accumulation signals using a 60× water immersion objective (1.1 NA, 0.2 WD) (Olympus, Hamburg, Germany). Cells were trapped in the laser focal point using optical trapping capability. At least 20 cells were randomly selected and measured in two distinct areas, one in the center and the other one near the cell membrane, in order to collect both cytoplasmic and nuclear spectral features.

### Raman spectra analysis

The Raman spectral data were processed and analyzed using the mpX-RamSES software (microphotonX GmbH, Tutzing, Germany; https://www.microphotonx.com/products-1). The Raman spectra were in a range of 480–1800 cm^−1^ containing the biological-relevant spectral bands. Baseline correction was also performed using an asymmetric least square fit, and the spectra were smoothened with a median filter. Next, the spectra were interpolated to continuous wave numbers and normalized by implementing unit vector normalization. The mpX-RamSES was also used for cellular and data visualization. Linear Discriminant Analysis (LDA) score plotwas then applied to illuminate the differences among the studied groups. Hierarchical cluster analysis (HCA) was applied as an unsupervised statistical analysis and clustering method to separate the Raman spectra based on the different spectral observations into different clusters.

### Endothelial cell migration assay

The HAECs were seeded 24 hours before treatment until they become confluent. Cell monolayer migration in response to a cell-free gap created by a sterile 200 μl pipet tip was assessed, as previously described. Images were continuously obtained at 6.4x magnification every 15 min from time point t=0 to t=16 h post-scratch using an incubator-equipped (humidified atmosphere, 37°C, 5% CO_2_) phase-contrast live cell imaging microscope (Olympus IX81) with the Hamamatsu (C11440) detector at 1-megapixel (1024*1024 pixel) 16 bit. The migrated area ratio at 16 h was measured using ImageJ software.

### Endothelial tube formation assay

The angiogenic capacity of endothelial cells represented as the number of tubes formed was examined in HAECs seeded onto Matrigel (Corning^®^ Matrigel^®^-356234) at the sub-confluent level. Images were continuously recorded at 6.4x magnification every 15 min from t=0 to t=16 h using an incubator-equipped phase-contrast Olympus IX81 microscope with the Hamamatsu (C11440) detector at 1-megapixel (1024*1024 pixel) 16 bit, and analyzed using ImageJ Angiogenesis Analyzer software.

### *Ex vivo* aortic ring sprouting assay

Aortic ring sprouting was assessed, as previously described. Fragments of murine aortas (2-mm) were cultured onto Matrigel and subsequently imaged daily (for 5 days) using an incubator-equipped phase-contrast Olympus IX81 microscope, followed by the analysis of the number of aortic sprouts.

### *Ex vivo* endothelium-dependent relaxation assay

The murine aortas at equal lengths (2☐mm) were cut and mounted in a 5-ml organ chamber filled with Krebs-Ringer bicarbonate solution (118.6☐mM NaCl, 4.7☐mM KCl, 2.5☐mM CaCl_2_, 1.2☐mM KH_2_PO_4_, 1.2☐mM MgSO_4_, 25.1☐mM NaHCO_3_, 2.6☐μM EDTA, 10.1☐mMglucose; 37☐°C, pH☐7.4), and bubbled with 95% O_2_, 5% CO_2_. Aortic rings were connected to an isometric force transducer (PowerLab 8/30 and LabChart v7.2.5, AD Instruments, Inc.) for continuous isometric tension recording (Multi-Myograph 610☐M, Danish Myo Technology, Denmark). Concentration–response curves were obtained in responses to increasing concentrations of acetylcholine (Ach, 10^−9^ to 10^−5^☐M; Sigma-Aldrich).

### Statistical analysis

Statistical differences were analyzed by unpaired two-tailed Student’s *t*-test, two-tailed Mann-Whitney *U*-test, and one-way or two-way Analysis of variance (ANOVA) followed by Tukey’s post *hoc* multiple comparison tests. At least three independent triplicated experiments were performed for each experimental set-up. Statistical analysis was performed using GraphPad Prism 8 software (v.8.0.1, La Jolla, CA, USA). The Raman spectra were analyzed using the mpX-RamSES software (microphotonX GmbH, Tutzing, Germany; https://www.microphotonx.com/products-1). Data are expressed as mean ± SD or SEM, and *P*<0.05 was defined as statistically significant, indicated as *P*<0.05 as *; *P*<0.01 as **; *P*<0.001 as ***; *P*<0.0001 as ****.

## Data availability

The authors declare that all data supporting the findings of this study are available within the paper and its Supplementary information files. The source data underlying Figs. 1B, 1D, 1F, 1G, 2B-D, 2F-H, 3A-E, 4B-G, and Supplemental Figs. 1B, 1C, 2B, 4B-D, and 5A are provided as the Source Data file.

All data from Raman spectroscopy analyses in this paper have been deposited at https://www.microphotonx.com/products-1 and are publicly available as of request. All other data and reagents that support the findings of this study are available from the lead contacts, Seyed Soheil Saeedi Saravi and Jürg H. Beer, upon request.

## Acknowledgement

We thank Prof. Thomas Michel (Division of Cardiovascular Medicine, Brigham and Women’s Hospital, Harvard Medical School) for helpful discussions and technical support (HyPer7.2 constructs).

The authors acknowledge funding from the Novartis Foundation for Medical-Biological Research (#21A053), the SwissLife Jubiläumsstiftung (#1286), the Stiftung Kardio, the Fonds zur Förderung des Akademischen Nachwuchses (FAN), and Gebauer Stiftung grants (to SSSS). JHB has been also funded by the Swiss National Science Foundation grant #310030_144152, Stiftung Kardio, and Swiss Heart Foundation.

## Contributions

SSSS conceptualized and designed the study. SSSS, KS, TS, KG, SL, and LP performed the experiments, including mouse studies, senescence and signal transduction studies, immunofluorescence assays, vasorelaxation studies, angiogenesis analyses, Seahorse mito stress assay, and single-cell micro-Raman spectroscopy. SSSS drafted and wrote the manuscript. SSSS, JHB, and FR contributed to scientific discussion. SSSS and JHB obtained the grant funding.

---

**Supplemental Fig. 1.**
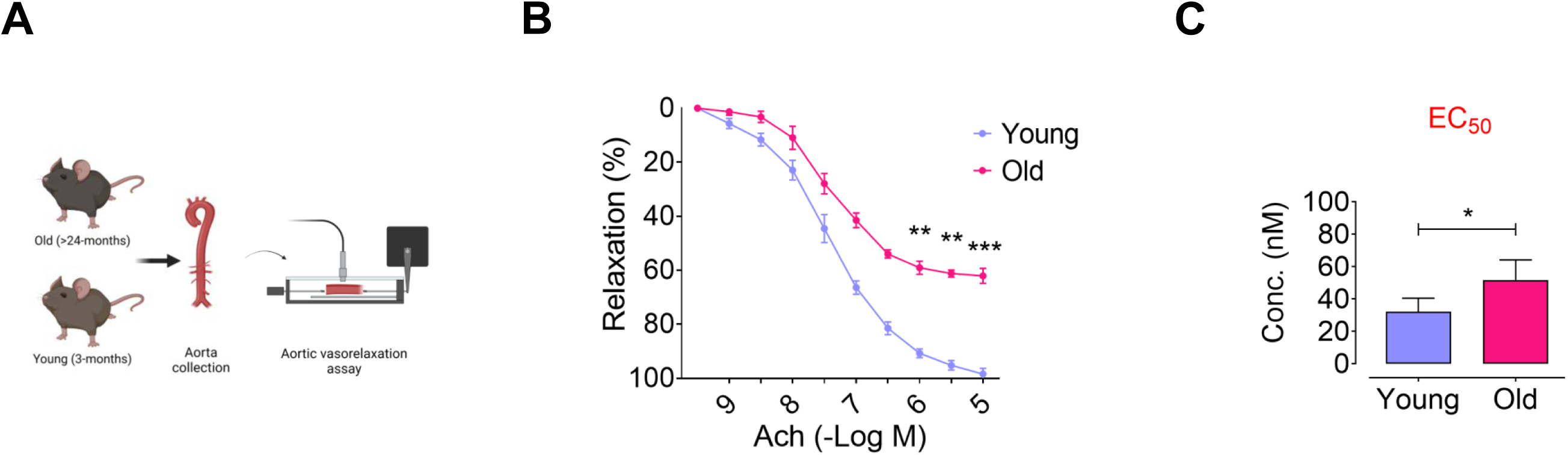
Aging is associated with endothelial dysfunction. **A**, Schematic diagram of the experimental setting: Aortas were harvested from old and young mice for *ex vivo* force tension myography in an organ bath setup. **B,** Vasorelaxation responses (%) *ex vivo* to acetylcholine (Ach), an endothelial-dependent mechanism, in aortic rings from old mice were significantly lower than young mice (n=10). **C,** Bar chart reveals a rightward shift of dose-response curve to Ach (higher EC_50_) in isolated aortas from old *versus* young mice. Continuous data are presented as mean ± SD. Statistical analysis was performed with a one-way ANOVA method (**B**) and two-tailed unpaired Student’s *t*-test (**C**). (\**P*<0.05, \*\**P*<0.01, \*\*\**P*<0.001). Source data are provided as a Source Data file.

**Supplemental Fig. 2.**
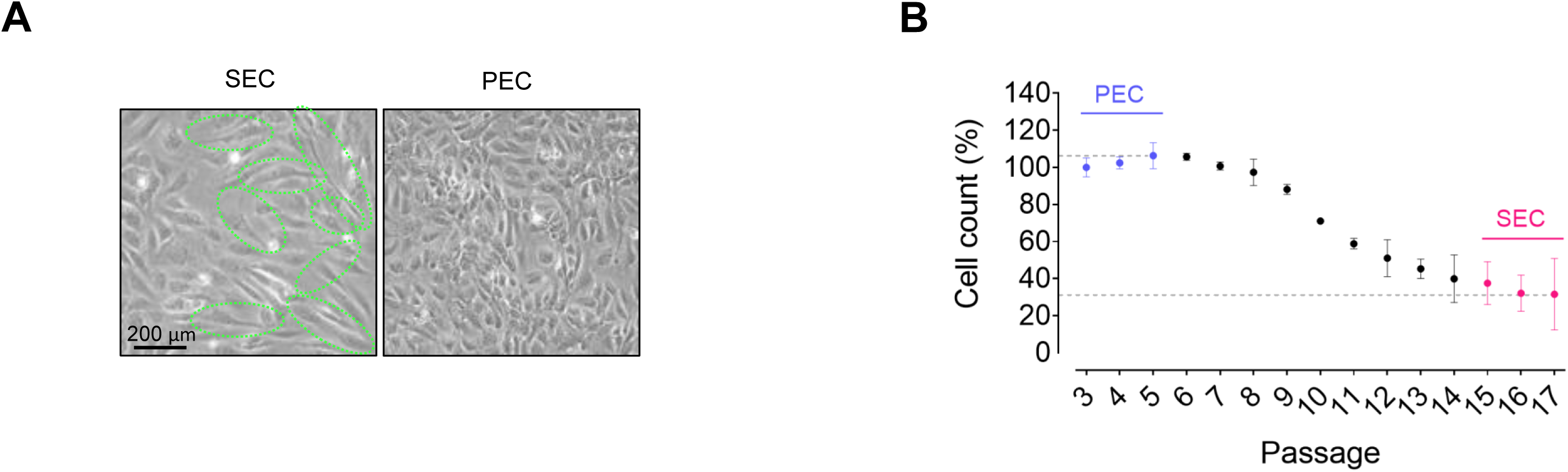
Replicative senescence phenotype in endothelial cells. **A**, Representative bright-field images depict cellular senescence-like phenotype, including enlarged, flattened, and multinucleated appearance, in replicative SECs. The images demonstrate no morphologically senescence-like phenotype in PECs (n=6). **B,** HAEC were counted at each passage from p.3 to p.17 and reported as percentage (%). The cells at passages 3 to 5 were characterized as PEC, while those at passages 15 to 17 were considered as SEC (n=6). Scale bar, 200 μm. Green circles indicate senescent-like cells. Experiments were triplicated independently.

**Supplemental Fig. 3.**
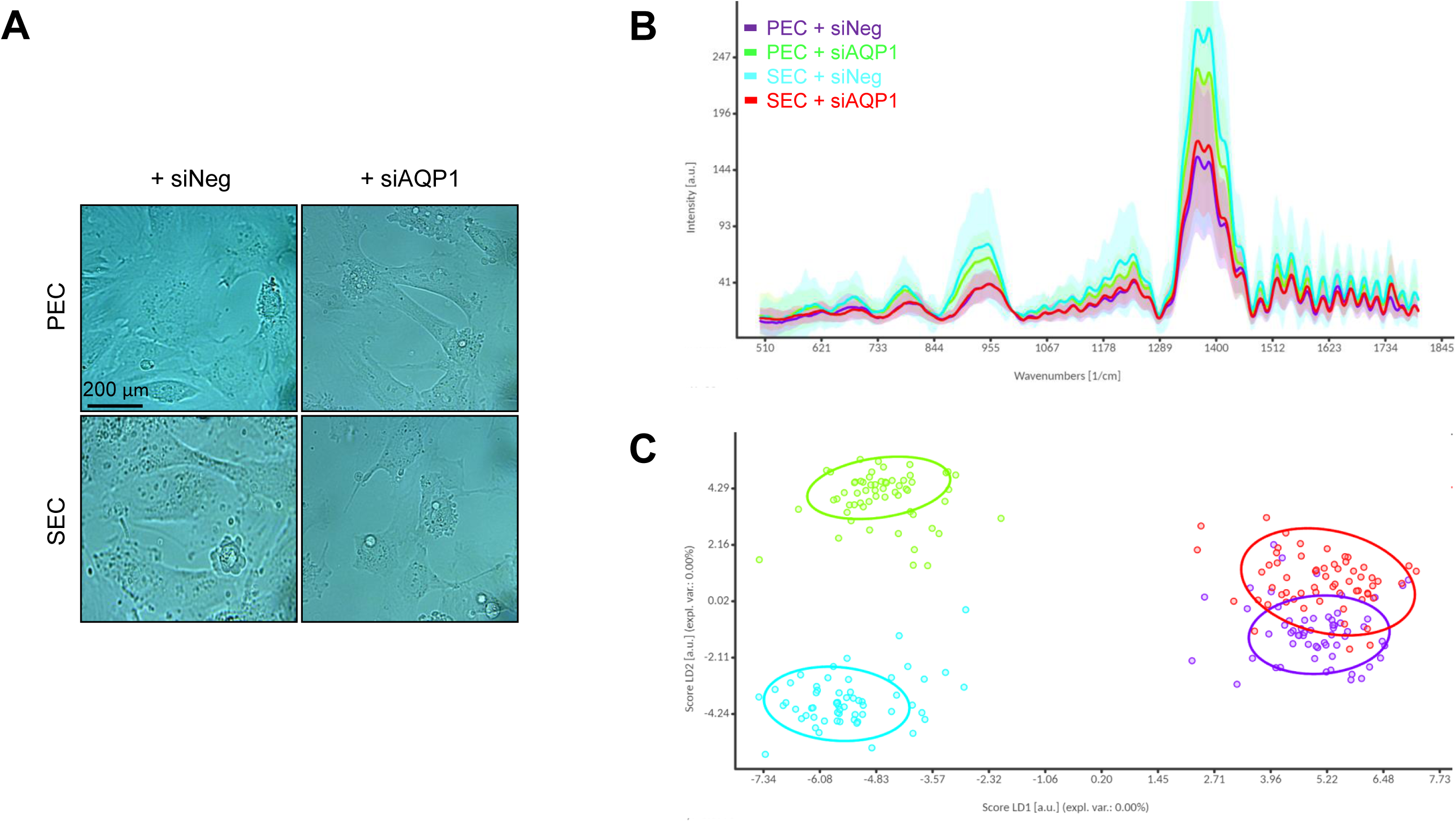
AQP1 differentially mediates endothelial secretome. **A**, Representative bright-field images of siNeg-or siAQP1-transfected PECs *versus* SECs cultured in multichannel μ-slides. **B,** Single-cell Raman spectra reveals relatively distinct phenotypical resemblance of PECs and SECs transfected with either siNeg or siAQP1 in a 2D culture (n=100). The peaks were vector-normalized. **C,** Linear Discriminant Analysis (LDA) score plot shows four distinct populations of siNeg-PECs (violet dots), siAQP1-PECs (green dots), siNeg-SECs (cyan dots), and siAQP1-SECs (red dots). Each single dot represents a Raman spectral readout, and the ellipses represent 90% CI. Scale bar, 20 μm.

**Supplemental Fig. 4.**
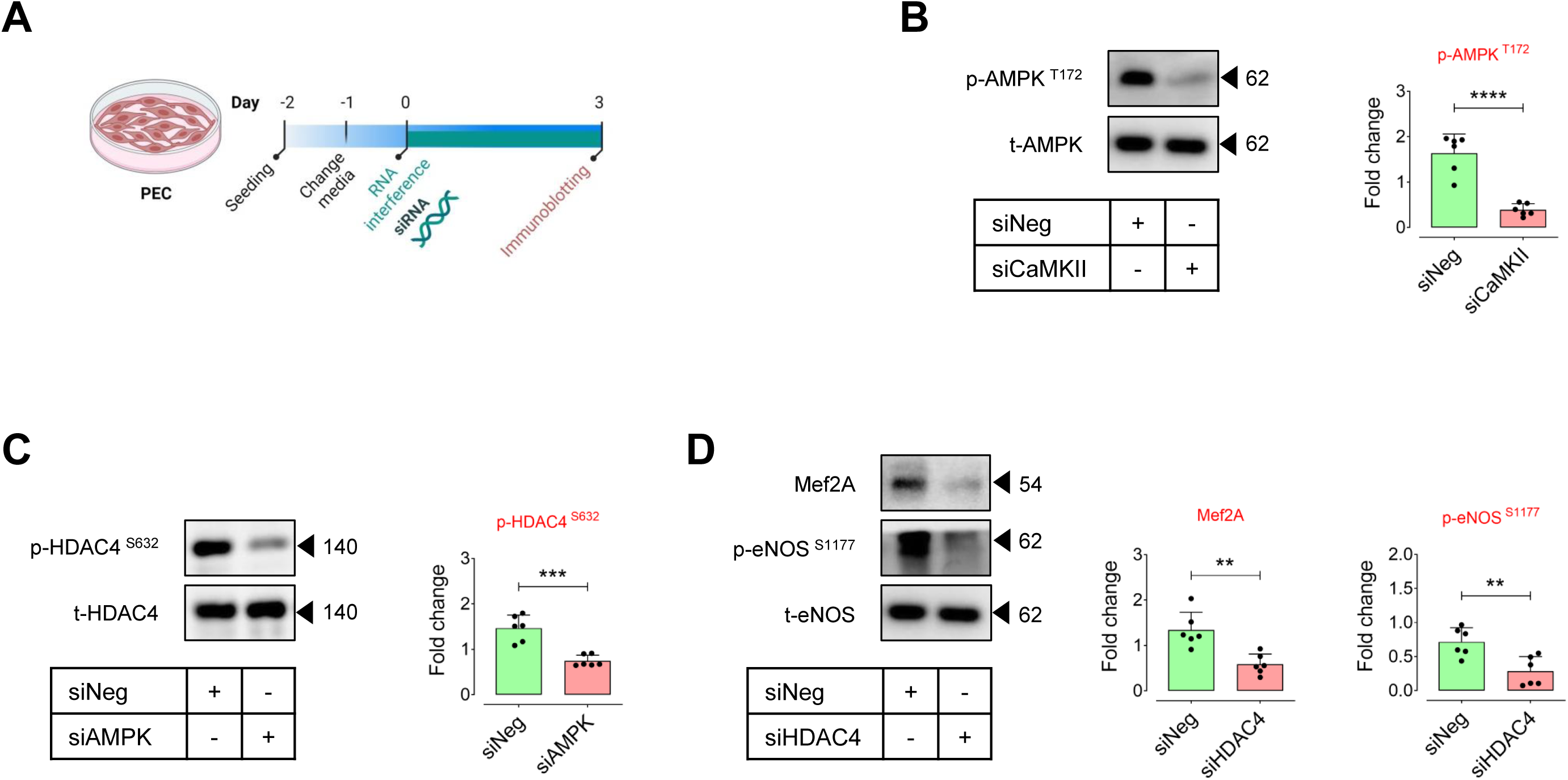
Exogenous H_2_O_2_ regulates endothelial function through CaMKII-HDAC4 pathway. **A**, Schematic diagram of the experimental setting: PECs were transfected with siCaMKII, siAMPK, or siHDAC4 in the presence of H_2_O_2_ (25 μM). **B-D,** Immunoblots (left) and quantitative plots (right) characterizing the effects of exogenous H_2_O_2_ on post-translational modifications of AMPK (**B**) and its downstream epigenetic regulator HDAC4 (**C**) by CaMKII and AMPK knockdown, respectively, in PECs (n=6). **D,** Analysis of Mef2A expression and eNOS phosphorylation at Ser1177 in siHDAC4-transfected PECs (n=6). Error bars represent SD (**B-D**). Data represent triplicated biologically independent experiments. *P* values were calculated using two-tailed unpaired Student’s *t*-test (**B-D**). (\*\**P*<0.01, \*\*\**P*<0.001, \*\*\*\**P*<0.0001). Source data are provided as a Source Data file.

**Supplemental Fig. 5.**
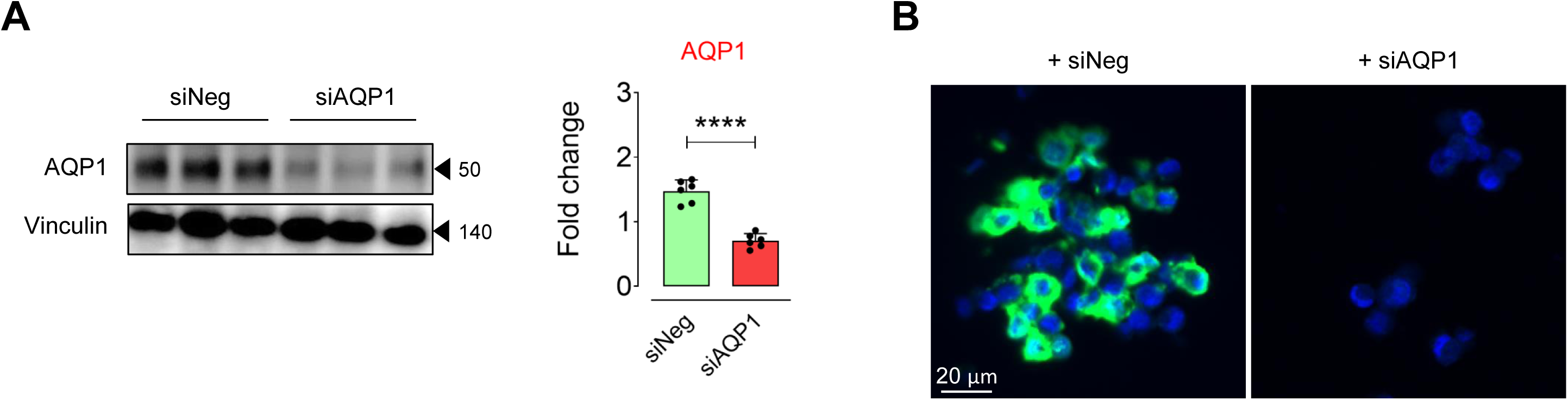
Characterization of siAQP1 in ECs. **A,B**, Immunoblots (**A**) and immunostaining (**B**) represent a marked reduction in the AQP1 expression in PECs transfected with siAQP1 (n=6). Scale bar, 20 μm. Error bars represent SD (**A**). Data represent triplicated biologically independent experiments. *P* value was calculated using two-tailed unpaired Student’s *t*-test (**A**). (\*\*\*\**P*<0.0001). Source data are provided as a Source Data file.

**Supplemental Table 1.**
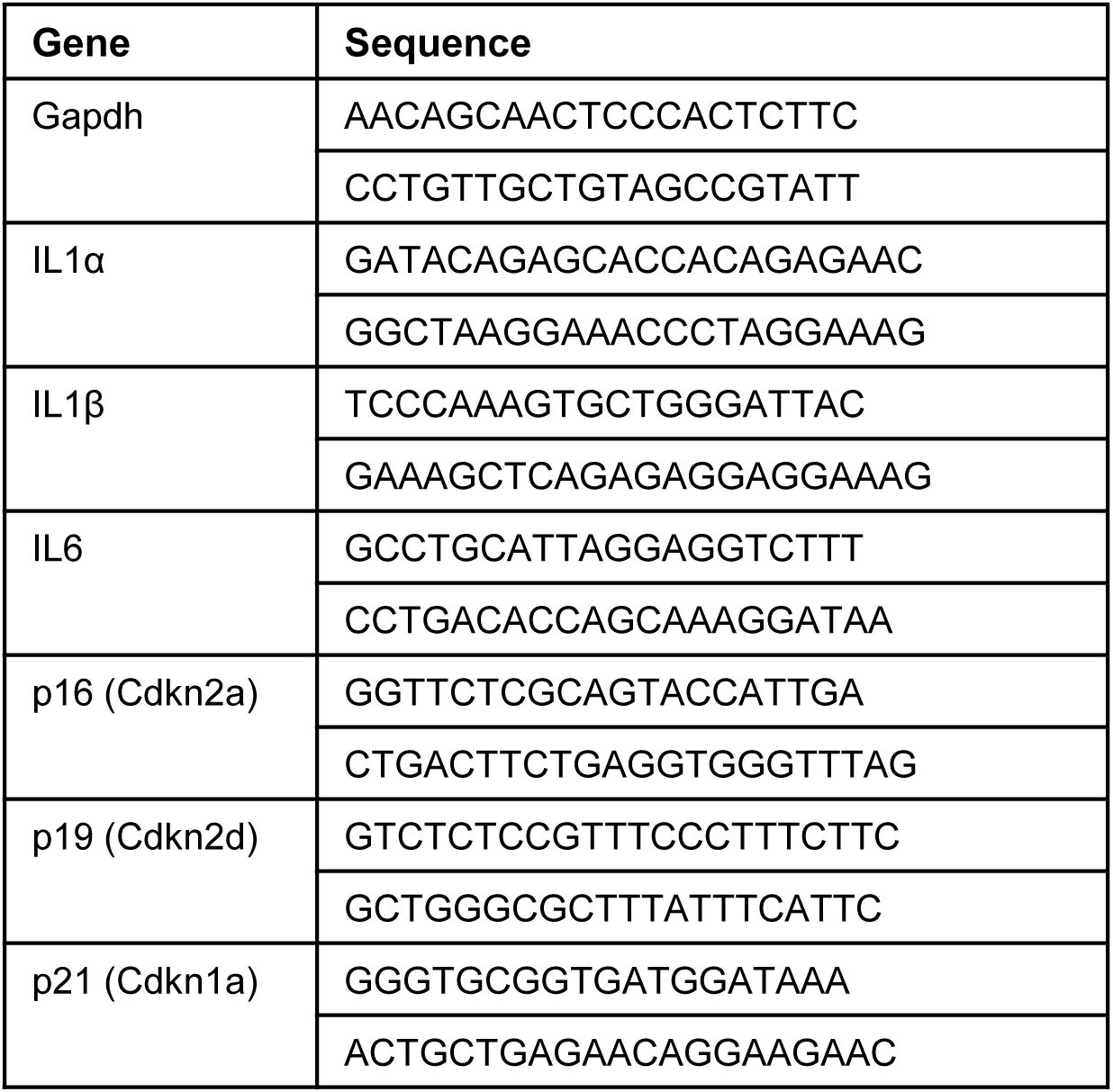
Nucleotide sequence of primers used in this study.

## References

1. Bloom, S. I. et al. Mechanisms and consequences of endothelial cell senescence. Nat. Rev. Cardiol. 20, 38–51 (2023).

2. Allemann, M. S. et al. Targeting the redox system for cardiovascular regeneration in aging. Aging Cell. 22, e14020 (2023).

3. Gasek, N. S. et al. Strategies for targeting senescent cells in human disease. *Nat*. Aging. 1, 870–879 (2021).

4. Han, Y. & Kim, S. Y. Endothelial senescence in vascular diseases: current understanding and future opportunities in senotherapeutics. Exp. Mol. Med. 55, 1–12 (2023).

5. Yrrell, D. J. & Goldstein, D. R. Ageing and atherosclerosis: vascular intrinsic and extrinsic factors and potential role of IL-6. Nat. Rev. Cardiol. 18, 58–68 (2021).

6. Saeedi Saravi, S. S. & Feinberg, M.W. Can removal of zombie cells revitalize the aging cardiovascular system? Eur. Heart J. ehad849 (2024).

7. Donato, A. J. et al. Mechanisms of dysfunction in the aging vasculature and role in age-related disease. Circ. Res. 123, 825–848 (2018).

8. Stabenow, L. K. et al. Oxidative glucose metabolism promotes senescence in vascular endothelial cells. Cells. 11, 2213 (2022).

9. Ww

10. Saeedi Saravi, S.S., et al. Differential endothelial signaling responses elicited by chemogenetic H_2_O_2_ synthesis. Redox Biol. 36, 101605 (2020).

11. Miller, A. W. et al. Aquaporin-3 mediates hydrogen peroxide uptake to regulate downstream intracellular signaling. Proc. Natl. Acad. Sci. USA. 107, 15681–15686 (2010).

12. Cai, H. Hydrogen peroxide regulation of endothelial function: origins, mechanisms, and consequences. Cardiovasc. Res. 68, 26–36 (2005).

13. Tomalin, L. E. et al. Increasing extracellular H_2_O_2_ produces a bi-phasic response in intracellular H_2_O_2_, with peroxiredoxin hyperoxidation only triggered once the cellular H_2_O_2_-buffering capacity is overwhelmed. Free Radic. Biol. Med. 95, 333–348 (2016).

14. Montiel, V. et al. Inhibition of aquaporin-1 prevents myocardial remodeling by blocking the transmembrane transport of hydrogen peroxide. Sci. Transl. Med. 12, eaay2176 (2020).

15. Bienert, G. P. & Chaumont, F. Aquaporin-facilitated transmembrane diffusion of hydrogen peroxide. Biochim. Biophys. Acta 1840, 1596–1604 (2014).

16. Saadoun, S. et al. Impairment of angiogenesis and cell migration by targeted aquaporin-1 gene disruption. Nature. 434, 786–792 (2005).

17. Mun, G. I. et al. Differential gene expression in young and senescent endothelial cells under static and laminar shear stress conditions. Free Radic. Biol. Med. 47, 291–299 (2009).

18. Shangzu, Z. et al. Aquaporins: Important players in the cardiovascular pathophysiology. Pharmacol. Res. 183, 106363 (2022).

19. Bıçakçı, H. et al. Investigation of the effects of aging on the expression of aquaporin 1 and aquaporin 4 protein in heart tissue. Anatol. J. Cardiol. 17, 18–23 (2017).

20. Tammar, G. et al. Aquaporin Membrane Channels in Oxidative Stress, Cell Signaling, and Aging: Recent Advances and Research Trends. Oxid. Med. Cell. Longev. 2018, 1501847 (2018).

21. Han, Y. M. et al. β-Hydroxybutyrate Prevents Vascular Senescence through hnRNP A1-Mediated Upregulation of Oct4. Mol. Cell. 71, 1064–1078 (2018).

22. Saeedi Saravi, S. S., et al. Gut microbiota-dependent increase in phenylacetic acid induces endothelial cell senescence during aging. BioRxiv. 10.1101/2023.11.17.567594 (2023).

23. Chen, C. et al. Multiplexed live-cell profiling with Raman probes. Nat. Commun. 12, 3405 (2021).

24. Pontiggia, L., et al. Human Basal and Suprabasal Keratinocytes Are Both Able to Generate and Maintain Dermo–Epidermal Skin Substitutes in Long-Term In Vivo Experiments. Cells. 11, 2156 (2022).

25. Miwa, S. et al. Mitochondrial dysfunction in cell senescence and aging. J. Clin. Invest. 132, e158447 (2022).

26. Wu, Y. et al. Phosphoglycerate dehydrogenase activates PKM2 to phosphorylate histone H3T11 and attenuate cellular senescence. Nat. Commun. 14, 1323 (2023).

27. Sakamuri, S. S. V. P. et al. Glycolytic and oxidative phosphorylation defects precede the development of senescence in primary human brain microvascular endothelial cells. Geroscience. 44, 1975–1994 (2022).

28. Smith, J. K. & Johnson, A. B. Endothelial cell senescence and its implications for vascular remodeling. J. Vasc. Res. 45, 201–215 (2020).

29. Das, A. et al. Impairment of an endothelial NAD^+^-H_2_S signaling network is a reversible cause of vascular aging. Cell. 173, 74–89 (2018).

30. Kabacik, S. et al. The relationship between epigenetic age and the hallmarks of aging in human cells. *Nat*. Aging. 2, 484–493 (2022).

31. Wang, K. et al. Epigenetic regulation of aging: implications for interventions of aging and diseases. Sig. Transduct. Target Ther. 7, 374 (2022).

32. Schader, T. et al. Oxidation of HDAC4 by Nox4-derived H_2_O_2_ maintains tube formation by endothelial cells. Redox Biol. 36, 101669 (2020).

33. Lu, Y. W. et al. MEF2 (Myocyte Enhancer Factor 2) is essential for endothelial homeostasis and the atheroprotective gene expression program. Arterioscler. Thromb. Vasc. Biol. 41, 1105–1123 (2021).

34. Liu, B. et al. Myocyte enhancer factor 2A delays vascular endothelial cell senescence by activating the PI3K/p-Akt/SIRT1 pathway. Aging (Albany NY). 11, 3768–3784 (2019).

35. Eroglu, E. et al. Discordance between eNOS phosphorylation and activation revealed by multispectral imaging and chemogenetic methods. Proc. Natl. Acad. Sci. USA. 116, 20210–20217 (2019).

36. Huang, W. et al. Cellular senescence: the good, the bad and the unknown. Nat. Rev. Nephrol. 18, 611–627 (2022).

37. Agre, P. The aquaporin water channels. Proc. Am. Thorac. Soc. 3, 5–13 (2006).

38. Bienert, G. P. & Chaumont, F. Aquaporin-facilitated transmembrane diffusion of hydrogen peroxide. Biochim. Biophys. Acta. 1840, 1596–1604 (2014).

39. Lo Conte, M. & Carroll, K.S. The redox biochemistry of protein sulfenylation and sulfinylation. J. Biol. Chem. 288, 26480–26488 (2013).

40. Anasooya Shaji, C., et al. The Tri-phasic Role of Hydrogen Peroxide in Blood-Brain Barrier Endothelial cells. Sci. Rep. 9, 133 (2019).

41. Induction of endothelial NO synthase by hydrogen peroxide via a Ca2+/Calmodulin-dependent protein kinase II/Janus Kinase 2–dependent pathway. Arterioscler. Thromb. Vasc. Biol. 21, 1571–1576 (2001).

42. Maron, B. A. & Michel, T. Subcellular localization of oxidants and redox modulation of endothelial nitric oxide synthase. Circ. J. 76, 2497–2512 (2012).

43. Sies, H. Hydrogen peroxide as a central redox signaling molecule in physiological oxidative stress: Oxidative eustress. Redox Biol. 11, 613–619 (2017).

44. Sies, H. Oxidative eustress and oxidative distress: Introductory remarks. Academic Press, 3–12 (2020).

45. Zhuang, J. et al. Age-related accumulation of advanced oxidation protein products promotes osteoclastogenesis through disruption of redox homeostasis. Cell Death Dis. 12, 1160 (2021).

46. Chen, Y. et al. p53 SUMOylation Mediates AOPP-Induced Endothelial Senescence and Apoptosis Evasion. Front. Cardiovasc. Med. 8, 795747 (2022).

47. Li, L. et al. Nintedanib ameliorates oxidized low-density lipoprotein-induced inflammation and cellular senescence in vascular endothelial cells. Bioengineered. 13, 6196–6207 (2022).

48. Gomez-Cabrera, M. C. et al. Oxidative stress and mitochondrial impairment in aging. Free Radic. Biol. Med. 65, 26–37 (2016).

49. Victorelli, S. et al. Apoptotic stress causes mtDNA release during senescence and drives the SASP. Nature. 622, 627–636 (2023).

50. Baker, D. J. et al. Clearance of p16^Ink4a^-positive senescent cells delays ageing-associated disorders. Nature. 479, 232–236 (2011).

51. Fonseca, C. G. et al. Endothelial cells on the move: dynamics in vascular morphogenesis and disease. *Vasc*. Biol. 2, H29–H43 (2020).

52. Chen, X. et al. Therapeutic angiogenesis and tissue revascularization in ischemic vascular disease. J. Biol. Eng. 17, 13 (2023).

53. Matsushima, S. et al. Increased oxidative stress in the nucleus caused by Nox4 mediates oxidation of HDAC4 and cardiac hypertrophy. Circ. Res. 112, 651–663 (2013).

54. Fukumura, D. et al. Predominant role of endothelial nitric oxide synthase in vascular endothelial growth factor-induced angiogenesis and vascular permeability. Proc. Natl. Acad. Sci. USA. 98, 2604–2609 (2001).

55. Chacar, S. et al. Role of CaMKII in diabetes induced vascular injury and its interaction with anti-diabetes therapy. Rev. Endocr. Metab. Disord. (2023). 10.1007/s11154-023-09855-9.

56. Zheng, Z. et al. Ginsenoside Rb1 reduces H2O2-induced HUVEC dysfunction by stimulating the sirtuin-1/AMP-activated protein kinase pathway. Mol. Med. Rep. 22, 247–256 (2020).

57. Han, X. et al. AMPK activation protects cells from oxidative stress-induced senescence via autophagic flux restoration and intracellular NAD(+) elevation. Aging Cell. 15, 416–427 (2016).

58. Smith, T.L. et al. eNOS controls angiogenic sprouting and retinal neovascularization through the regulation of endothelial cell polarity. Cell. Mol. Life Sci. 79, 37 (2022).

59. Sawada, N. et al. Regulation of endothelial nitric oxide synthase and postnatal angiogenesis by Rac1. Circ. Res. 103, 360–368 (2008).

60. Spyropoulos, F. et al. Metabolomic and transcriptomic signatures of Chemogenetic heart failure. Am. J. Physiol. Heart Circ. Physiol. 322, H451–H465 (2021).

61. Sorrentino, A. et al. Reversal of heart failure in a chemogenetic model of persistent cardiac redox stress. Am. J. Physiol. Heart Circ. Physiol. 317, H617–H626 (2019).

